# Current clinical use of polygenic scores will risk exacerbating health disparities

**DOI:** 10.1101/441261

**Authors:** Alicia R. Martin, Masahiro Kanai, Yoichiro Kamatani, Yukinori Okada, Benjamin M. Neale, Mark J. Daly

## Abstract

Polygenic risk scores (PRS) are poised to improve biomedical outcomes via precision medicine. However, the major *ethical* and *scientific* challenge surrounding clinical implementation is that they are many-fold more accurate in European ancestry individuals than others. This disparity is an inescapable consequence of Eurocentric genome-wide association study biases. This highlights that—unlike clinical biomarkers and prescription drugs, which may individually work better in some populations but do not ubiquitously perform far better in European populations—clinical uses of PRS today would systematically afford greater improvement to European descent populations. Early diversifying efforts show promise in levelling this vast imbalance, even when non-European sample sizes are considerably smaller than the largest studies to date. To realize the full and equitable potential of PRS, we must prioritize greater diversity in genetic studies and public dissemination of summary statistics to ensure that health disparities are not increased for those already most underserved.

---

Polygenic risk scores (PRS), which predict complex traits using genetic data, are of burgeoning interest to the clinical community as researchers demonstrate their growing power to improve clinical care, genetic studies of a wide range of phenotypes increase in size and power, and genotyping costs plummet to less than US$50. Many earlier criticisms of limited prediction power are now recognized to have been chiefly an issue of insufficient sample size, which is no longer the case for many outcomes^1^. For example, polygenic risk scores alone already predict breast cancer, prostate cancer, and type 1 diabetes risk in European descent patients more accurately than current clinical models^2–4^. Additionally, integrated models of PRS together with other lifestyle and clinical factors have enabled clinicians to more accurately quantify the risk of heart attack for patients; consequently, they have more effectively targeted the reduction of LDL cholesterol and by extension heart attack by prescribing statins to patients at the greatest overall risk of cardiovascular disease^5–9^. Promisingly, return of genetic risk of complex disease to at-risk patients does not induce significant self-reported negative behavior or psychological function, and some potentially positive behavioral changes have been detected^10^. While we share enthusiasm about the potential of PRS to improve health outcomes through their eventual routine implementation as clinical biomarkers, we consider the consistent observation that they are currently of far greater predictive value in individuals of recent European descent than in others to be the major *ethical* and *scientific* challenge surrounding clinical translation and, at present, the most critical limitation to genetics in precision medicine. The scientific basis of this imbalance has been demonstrated theoretically, in simulations, and empirically across many traits and diseases^11–22^.

All studies to date using well-powered genome-wide association studies (GWAS) to assess the predictive value of PRS across a range of traits and populations have made a consistent observation: PRS predict individual risk far more accurately in Europeans than non-Europeans^15,16,18–24^. Rather than chance or biology, this is a predictable consequence of the fact that the genetic discovery efforts to date heavily underrepresent non-European populations globally. The correlation between true and genetically predicted phenotypes decays with genetic divergence from the makeup of the discovery GWAS, meaning that the accuracy of polygenic scores in different populations is highly dependent on the study population representation in the largest existing ‘training’ GWAS. Here, we document study biases that underrepresent non-European populations in current GWAS, and explain the fundamental concepts contributing to reduced phenotypic variance explained with increasing genetic divergence from populations included in GWAS.

## Predictable basis of disparities in PRS accuracy

Poor generalizability of genetic studies across populations arises from the overwhelming abundance of European descent studies and dearth of well-powered studies in globally diverse populations^25–28^. According to the GWAS catalog, ∼79% of all GWAS participants are of European descent despite making up only 16% of the global population (**Figure 1**). This is especially problematic as previous studies have shown that Hispanic/Latino and African American studies contribute an outsized number of associations relative to studies of similar sizes in Europeans^27^. More concerningly, the fraction of non-European individuals in GWAS has stagnated or declined since late 2014 (**Figure 1**), suggesting that we are not on a trajectory to correct this imbalance. These numbers provide a composite metric of study availability, accessibility, and use— cohorts that have been included in numerous GWAS are represented multiple times, which may disproportionately include cohorts of European descent. However, whereas the average sample sizes of GWAS in Europeans continue to grow, they have stagnated and remain several-fold smaller in other populations (**Supplementary Figure 1**).

**Figure 1.**
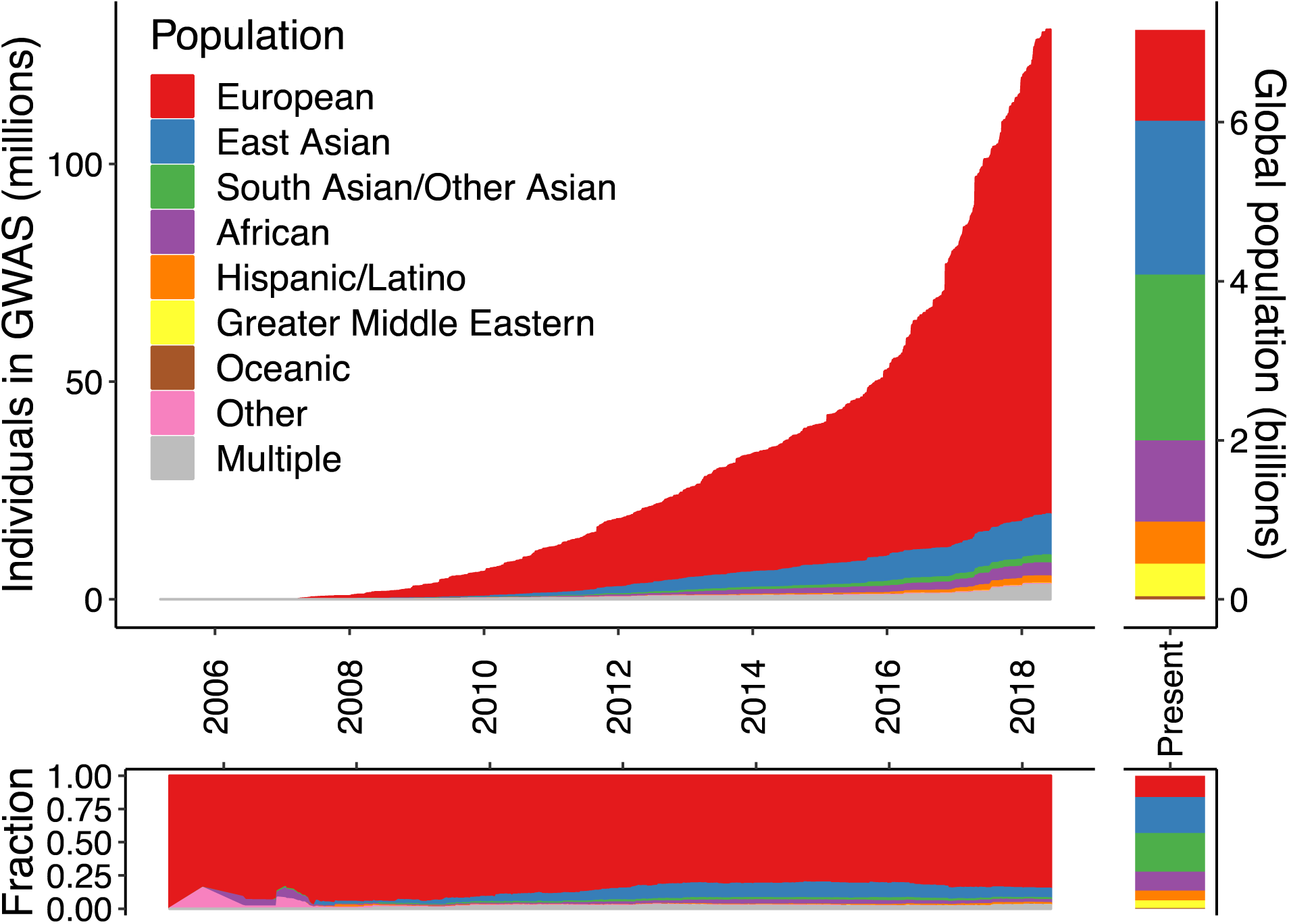
Ancestry of GWAS participants over time compared to the global population. Cumulative data as reported by the GWAS catalog^76^. Individuals whose ancestry is “not reported” are not shown.

The relative sample compositions of GWAS result in highly predictable disparities in prediction accuracy; population genetics theory predicts that genetic risk prediction accuracy will decay with increasing genetic divergence between the original GWAS sample and target of prediction, a function of population history^13,14^. This pattern can be attributed to several statistical observations which we detail below: 1) GWAS favor the discovery of genetic variants that are common in the study population; 2) linkage disequilibrium (LD) differentiates marginal effect size estimates for polygenic traits across populations, even when causal variants are the same; and 3) environment and demography differ across populations. Notably, the first two phenomena degrade prediction performance across populations substantially even when there exist no biological, environmental, or diagnostic differences, whereas the environment and demography may interact to drive differential forces of natural selection that in turn drive differences in causal genetic architecture. (We define the causal genetic architecture as the true effects of variants that impact a phenotype that would be identified in a population of infinite sample size. Unlike effect size estimates, true effects are typically modeled as invariant with respect to LD and allele frequency differences across populations.)

### Common discoveries and low-hanging fruit

First, the power to discover an association in a genetic study depends on the effect size and frequency of the variant^29^. This dependence means that the most significant associations tend to be more common in the populations in which they are discovered than elsewhere^13,30^. For example, GWAS catalog variants are more common on average in European populations compared to East Asian and African populations (**Figure 2B**), an observation not representative of genomic variants at large. Understudied populations offer low-hanging fruit for genetic discovery because variants that are common in these groups but rare or absent in European populations could not be discovered even with very large European sample sizes. Some examples include *SLC16A11* and *HNF1A* associations with type II diabetes in Latino populations, as well as *APOL1* associations with end-stage kidney disease and associations with prostate cancer in African descent populations^31–34^. If we assume that causal genetic variants have an equal effect across all populations—an assumption with some empirical support that offers the best case scenario for transferability^35–40^—Eurocentric GWAS biases mean that variants associated with risk are disproportionately common and discovered in European populations, accounting for a larger fraction of the phenotypic variance there^13^. Furthermore, imputation reference panels share the same study biases as in GWAS^41^, creating challenges for imputing sites that are rare in European populations but common elsewhere when the catalog of non-European haplotypes is substantially smaller. These issues are insurmountable through statistical methods alone^13^, but rather motivate substantial investments in more diverse populations to produce similar-sized GWAS of biomedical phenotypes in other populations.

**Figure 2.**
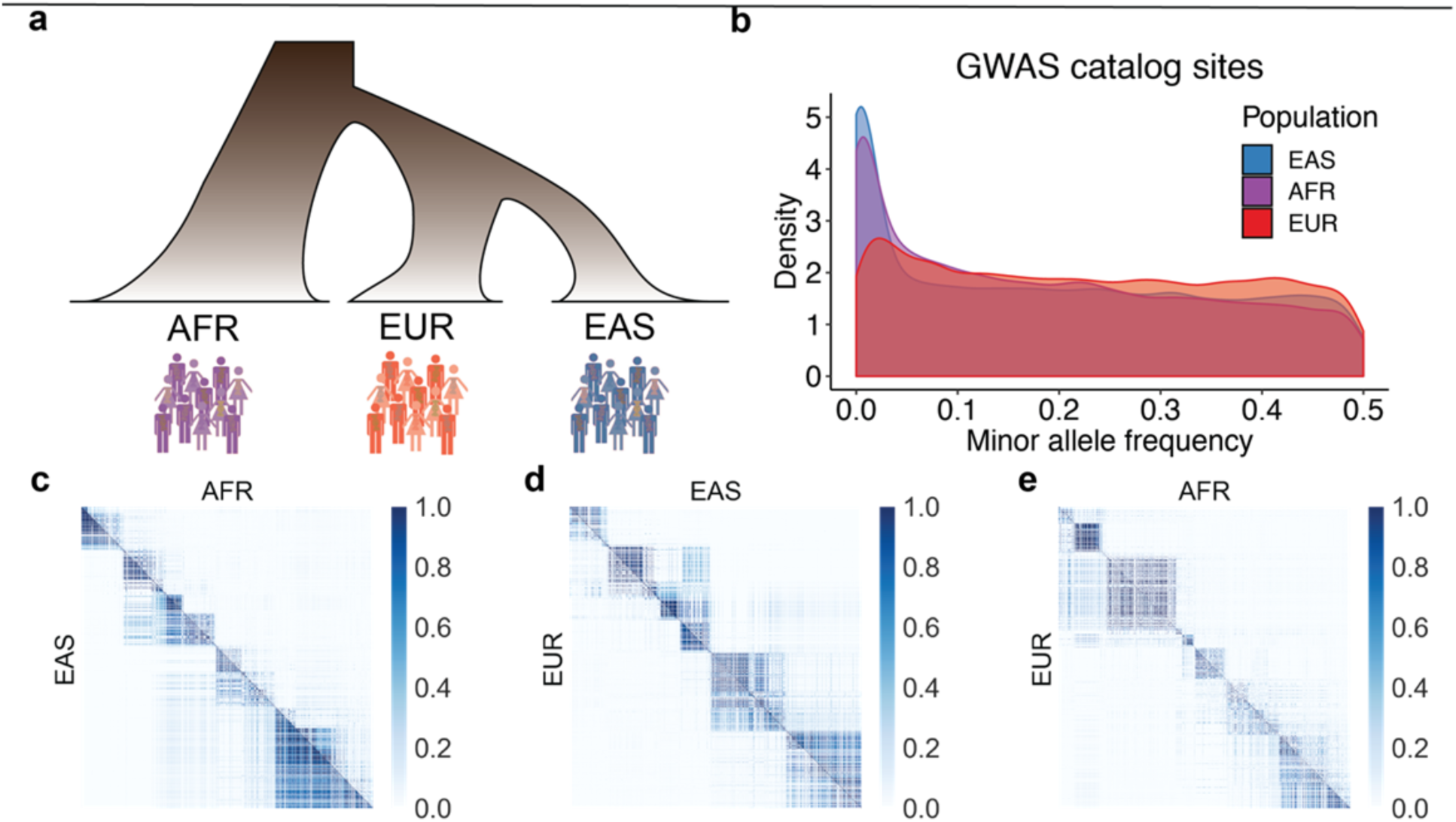
Demographic relationships, allele frequency differences, and local LD patterns between population pairs. Data analyzed from 1000 Genomes, in which population labels are: AFR = continental African, EUR = European, and EAS = East Asian. **a**) Cartoon relationships among AFR, EUR, and EAS populations. **b**) Allele frequency distributions in AFR, EUR, and EAS populations of variants from the GWAS catalog. **c-e**) Color axis shows LD scale (*r^2^*). LD comparisons between pairs of populations show the same region of the genome for each comparison (representative region is chr1, 51572kb-52857 kb) among pairs of SNPs polymorphic in both populations, illustrating that different SNPs are polymorphic across some population pairs, and that these SNPs have variable LD patterns across populations.

### Linkage disequilibrium

Second, LD, the correlation structure of the genome, varies across populations due to demographic history (**Figure 2A,C-E**). These LD differences in turn drive differences in effect size estimates (i.e. predictors) from GWAS across populations in proportion to LD between tagging and causal SNP pairs, even when causal effects are the same ^35,37–40^ (**Supplementary Note**). Differences in effect size estimates due to LD differences may typically be small for most regions of the genome (**Figure 2C-E**), but PRS sum across these effects, also aggregating these population differences. While it would be ideal to use causal effects rather than correlated effect size estimates to calculate PRS, it may not be feasible to fine-map most variants to a single locus to solve issues of low generalizability, even with very large GWAS. This is because complex traits are highly polygenic, meaning most of our prediction power comes from small effects that do not meet genome-wide significance and/or cannot be fine-mapped, even in many of the best-powered GWAS to date^42^.

### Complexities of history, selection, and the environment

Lastly, other cohort considerations may further worsen prediction accuracy differences across populations in less predictable ways. GWAS ancestry study biases and LD differences across populations are extremely challenging to address, but these issues actually make many favorable assumptions that all causal loci have the same impact and are under equivalent selective pressure in all populations. In contrast, other effects on polygenic adaptation or risk scores such as long-standing environmental differences across global populations that have resulted in differing responses of natural selection can impact populations differently based on their unique histories. Additionally, residual uncorrected population stratification may impact risk prediction accuracy across populations, but the magnitude of its effect is currently unclear. These effects are particularly challenging to disentangle, as has clearly been demonstrated for height, where evidence of polygenic adaptation and/or its relative magnitude is under question^43,44^. Comparisons of geographically stratified phenotypes like height across populations with highly divergent genetic backgrounds and mean environmental differences, such as differences in resource abundance during development across continents, are especially prone to confounding from correlated environmental and genetic divergence^43,44^. This residual stratification can lead to over-predicted differences across geographical space^45^.

Related to stratification, most PRS methods do not explicitly address recent admixture and none consider recently admixed individuals’ unique local mosaic of ancestry; further methods development is needed. Additionally, comparing PRS across environmentally stratified cohorts, such as in some biobanks with healthy volunteer effects versus disease study datasets or hospital-based cohorts, requires careful consideration of technical differences, collider bias, as well as variability in baseline health status among studies. It is also important to consider differences in definitions of clinical phenotypes and heterogeneity of sub-phenotypes among countries.

Differences in environmental exposure, gene-gene interactions, gene-environment interactions, historical population size dynamics, statistical noise, some potential causal effect differences, and/or other factors will further limit generalizability for genetic risk scores in an unpredictable, trait-specific fashion^46–49^. Complex traits do not behave in a genetically deterministic manner, with some environmental factors dwarfing individual genetic effects, creating outsized issues of comparability across globally diverse populations. Among psychiatric disorders for example, whereas schizophrenia has a nearly identical genetic basis across East Asians and Europeans (*r_g_*=0.98)^40^, substantially different rates of alcohol use disorder across populations are partially explained by differences in availability and genetic differences impacting alcohol metabolism^50^. While non-linear genetic factors explain little variation in complex traits beyond a purely additive model^51^, some unrecognized nonlinearities and gene-gene interactions can also induce genetic risk prediction challenges, as pairwise interactions are likely to vary more across populations than individual SNPs. Mathematically, we can simplistically think of this in terms of a two-SNP model, in which the sum of two SNP effects is likely to explain more phenotypic variance than the product of the same SNPs. Some machine learning approaches may thus modestly improve PRS accuracy beyond current approaches for some phenotypes^52^, but most likely for atypical traits with simpler architectures, known interactions, and poor prediction generalizability across populations, such as skin pigmentation^53^.

## Limited generalizability of PRS across diverse populations

So far, multi-ethnic work has been slow in most disease areas^54^, limiting even the opportunity to assess PRS in non-European cohorts. Nonetheless, some previous work has assessed prediction accuracy across diverse populations in several traits and diseases for which GWAS summary statistics are available and identified large disparities across populations (**Supplementary Note**). These disparities are not simply methodological issues, as various approaches (e.g. pruning and thresholding versus LDPred) and accuracy metrics (R^2^ for quantitative traits and various pseudo-R^2^ metrics for binary traits) illustrate this consistently poorer performance in populations distinct from discovery samples across a range of polygenic traits (**Supplementary Table 1**). These assessments are becoming increasingly feasible with the growth and public availability of global biobanks as well as diversifying priorities from funding agencies^55,56^. We assessed how prediction accuracy decayed across globally diverse populations for 17 anthropometric and blood panel traits in the UK Biobank (UKBB) when using European-derived summary statistics (**Supplementary Note**). Consistent with previous studies, we find that relative to European prediction accuracy, genetic prediction accuracy was far lower in other populations: 1.6-fold lower in Hispanic/Latino Americans, 1.7-fold lower in South Asians, 2.5-fold lower in East Asians, and 4.9-fold lower in Africans on average (**Figure 3**).

**Figure 3.**
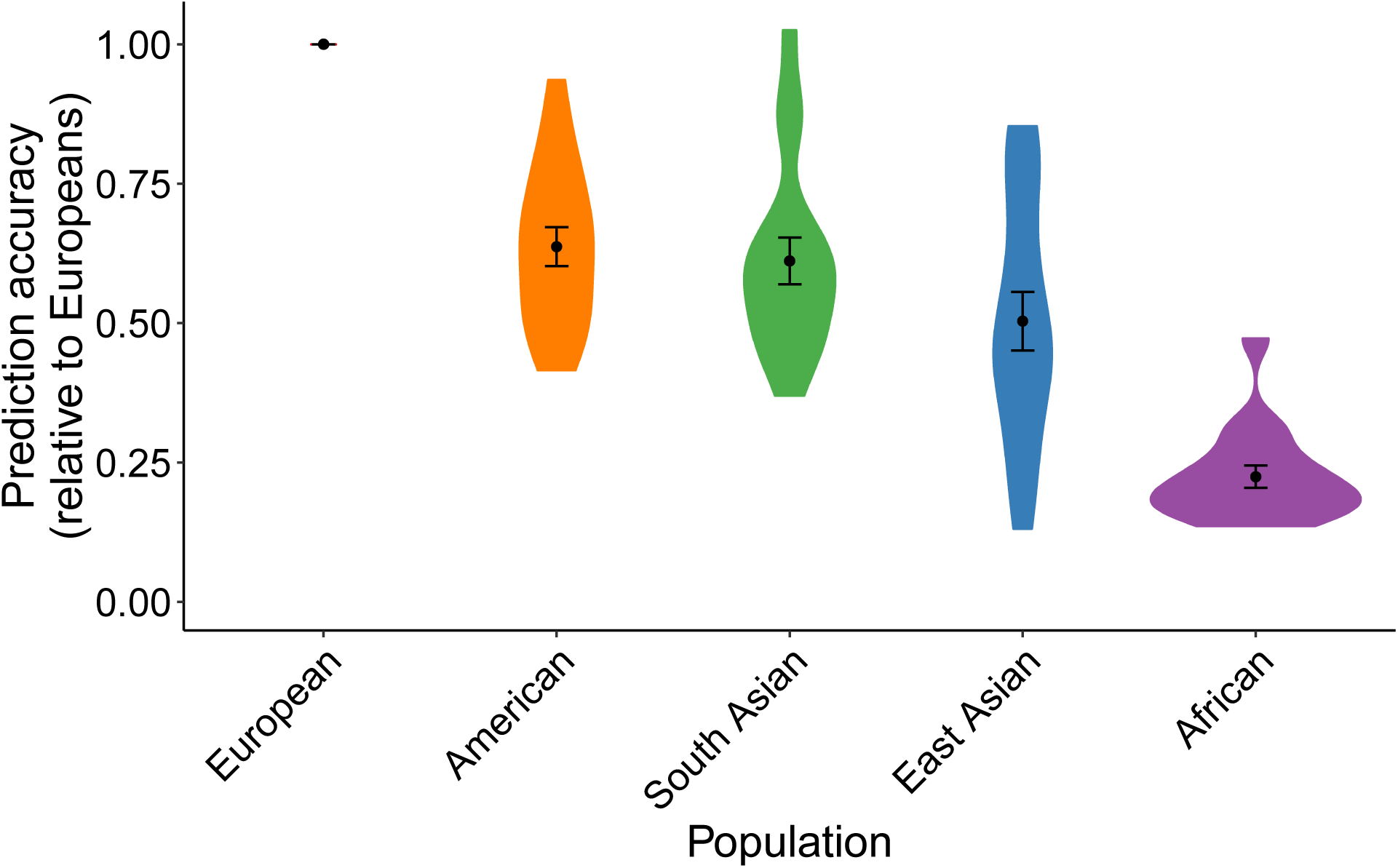
Prediction accuracy relative to European ancestry individuals across 17 quantitative traits and 5 continental populations in UKBB. All phenotypes shown here are quantitative anthropometric and blood panel traits, as described in **Supplementary Table 6**, which includes discovery cohort sample sizes. Prediction target individuals do not overlap with the discovery cohort and are unrelated, with sample sizes shown in **Supplementary Table 7**. Violin plots show distributions of relative prediction accuracies, points show mean values, and error bars show standard errors of the means. Prediction R^2^ for each trait and population are shown in **Supplementary Figure 12**.

## Prioritizing diversity shows early promise for PRS

Early diversifying GWAS efforts have been especially productive for informing on questions surrounding risk prediction. Rather than varying the prediction target dataset, some GWAS in diverse populations have increased the scale of non-European summary statistics and also varied the study dataset in multi-ethnic PRS studies^23,24,40^. These studies have shown that even when non-European cohorts are only a fraction the size of the largest European study, they are likely to have disproportionate value for predicting polygenic traits in other individuals of similar ancestry.

Given this background, we performed a systematic evaluation of polygenic prediction accuracy across 17 quantitative anthropometric and blood panel traits and five disease endpoints in British and Japanese individuals^23,57,58^ by performing GWAS with the exact same sample sizes in each population. We symmetrically demonstrate that prediction accuracy is consistently higher with GWAS summary statistics from ancestry-matched summary statistics (**Figure 4**, **Supplementary Figures 2-6**). Keeping in mind issues of comparability described above, we note that BBJ is a hospital-based disease-ascertained cohort, whereas UKBB is a healthier than average^59^ population-based cohort; thus, differences in observed heritability among these cohorts (rather than among populations) due to differences in phenotype precision likely explain lower prediction accuracy from the BBJ GWAS summary statistics for anthropometric and blood panel traits, but higher prediction accuracy for five ascertained diseases (**Supplementary Table 2**). Indeed, other East Asian studies have estimated higher heritability for some quantitative traits than BBJ using the same methods, such as for height (h^2^ = 0.48 ± 0.04 in Chinese women^60^). Some statistical fluctuations in the relative differences in prediction accuracy across populations are likely driven by differences in heritability measured in each population and/or trans-ethnic genetic correlation (i.e. of common variant effect sizes at SNPs common in two populations, **Supplementary Figures 7-10**, **Supplementary Tables 2–5**). These trans-ethnic correlation estimates indicate that effect sizes were mostly highly correlated across ancestries, with a few traits that were somewhat lower than excepted (e.g. height and BMI, with *ρ*_ge_=0.69 and 0.75, respectively). Prediction accuracy was far lower in individuals of African descent in the UK Biobank (**Supplementary Figures 4 and 11**) using GWAS summary statistics from either European or Japanese ancestry individuals, consistent with reduced prediction accuracy with increasing genetic divergence (**Figures 3 and 4**). These population studies demonstrate the power and utility of increasingly diverse GWAS for prediction, especially in populations of non-European descent.

**Figure 4.**
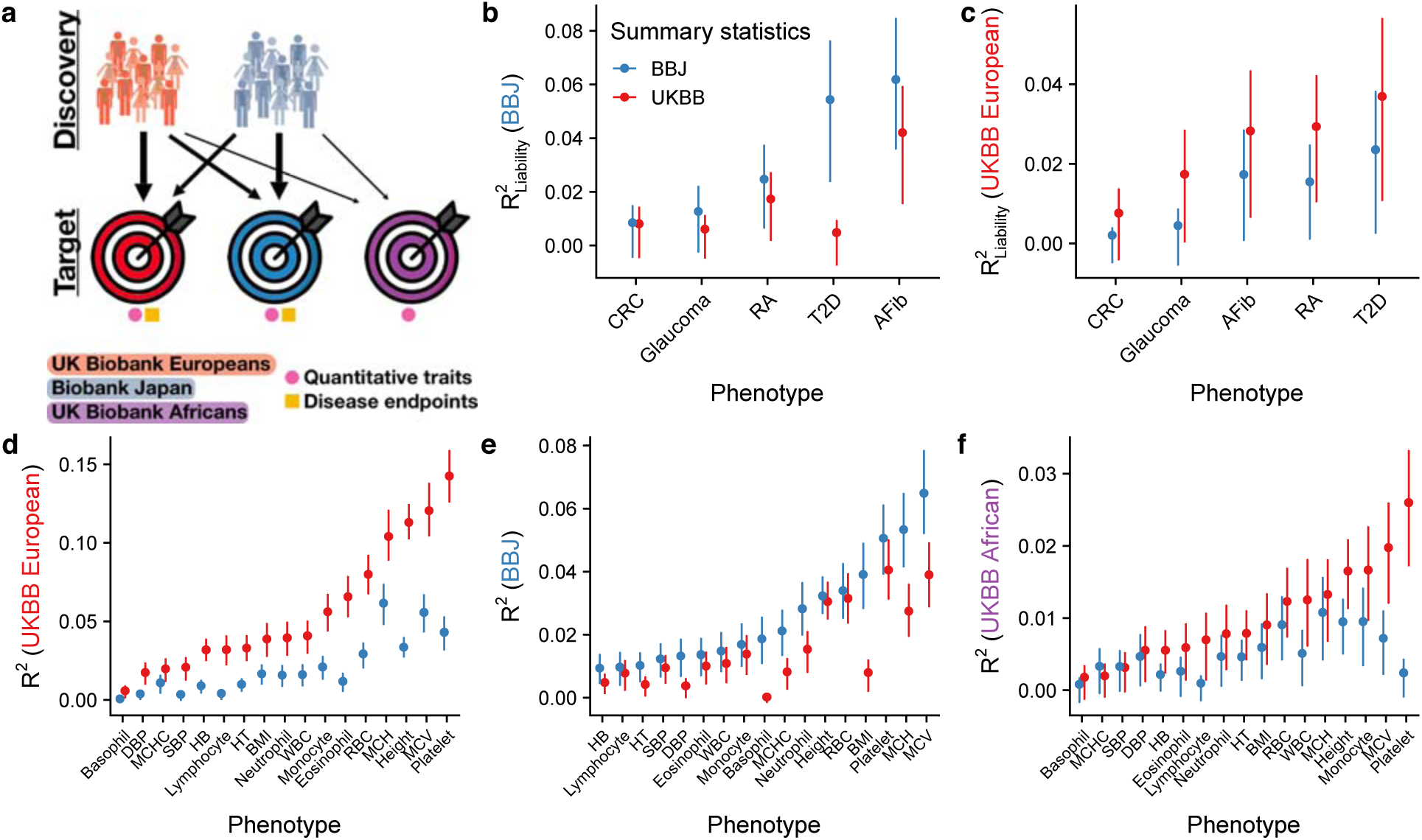
Polygenic risk prediction accuracy in Japanese, British, and African descent individuals using independent GWAS of equal sample sizes in the BioBank Japan (BBJ) and UK Biobank (UKBB). **a**) Explanatory diagram showing the different discovery and target cohorts/populations, and disease endpoints versus quantitative traits. **b-f**) Genetic prediction accuracy computed from independent BBJ and UKBB summary statistics with identical sample sizes (**Supplementary Tables 6 and 8**). Note that y-axes differ, reflecting differences in prediction accuracy. **b-c**) PRS accuracy for five diseases in: Japanese individuals in the BBJ (**b**) and British individuals in the UKBB. **d-f**) PRS accuracy for 17 anthropometric and blood panel traits in: Japanese individuals in the BBJ (**d**), British individuals in the UKBB (**e**), and African descent British individuals in the UKBB (**f**). Trait abbreviations are as in **Supplementary Table 6**. Each point shows the maximum R^2^ (i.e. best predictor) across five p-value thresholds, and lines correspond to 95% confidence intervals calculated via bootstrap. R^2^ values for all p-value thresholds tested are shown in **Supplementary Figures 2-6**. Prediction accuracy tends to be higher in the UKBB for quantitative traits than in BBJ and vice versa for disease endpoints, likely because of concomitant phenotype precision and consequently observed heritability for these classes of traits (**Supplementary Tables 2-4**). Thalassemia and sickle cell disease are unlikely to explain a significant fraction of prediction accuracy differences for blood panels across populations, as few individuals have been diagnosed with these disorders via ICD-10 codes (**Supplementary Table 9**).

While many other traits and diseases have been studied in multi-ethnic settings, few have reported comparable metrics of prediction accuracy across populations. Cardiovascular research, for example, has led the charge towards clinical translation of PRS^1^. This enthusiasm is driven by observations that a polygenic burden of LDL-increasing SNPs can confer monogenic-equivalent risk of cardiovascular disease, with polygenic scores improving clinical models for risk assessment and statin prescription that can reduce coronary heart disease and improve healthcare delivery efficiency^5–7^. However, many of these studies have been conducted exclusively in European descent populations, with few studies rigorously evaluating population-level applicability to non-Europeans. Those existing findings indeed demonstrate a large reduction in prediction utility in non-European populations^11^, though often with comparisons of odds ratios among arbitrary breakpoints in the risk distribution that make comparisons across studies challenging. To better clarify how polygenic prediction will be deployed in a clinical setting with diverse populations, more systematic and thorough evaluations of the utility of PRS within and across populations for many complex traits are still needed. These evaluations would benefit from rigorous polygenic prediction accuracy evaluations, especially for diverse non-European patients^61–63^.

## Clinical use of PRS may uniquely exacerbate disparities

Our impetus for raising these statistical issues limiting the generalizability of PRS across population stems from our concerns that, while they are legitimately clinically promising for improving health outcomes for many biomedical phenotypes, they may have a larger potential to raise health disparities than other clinical factors for several reasons. The opportunities they provide for improving health outcomes means they inevitably will and should be pursued in the near term, but we urge that a concerted prioritization to make GWAS summary statistics easily accessible for diverse populations and a variety of traits and diseases is imperative, even when they are a fraction the size of the largest existing European datasets. *Individual* clinical tests, biomarkers, and prescription drug efficacy may vary across populations in their utility, but are fundamentally informed by the same underlying biology^64,65^. Currently, guidelines state that as few as 120 individuals define reference intervals for clinical factors (though often smaller numbers from only one subpopulation are used) and there is no clear definition of who is “normal”^64^. Consequently, reference intervals for biomarkers can sometimes deviate considerably by reported ethnicity^66–68^. Defining ethnicity-specific reference intervals is clearly an important problem that can provide fundamental interpretability gains with implications for some major health benefits (e.g. need for dialysis and development of Type 2 diabetes based on ethnicity-specific serum creatinine and hemoglobin A1C reference intervals, respectively)^67^. Simply put, some biomarkers or clinical tests scale directly with health outcomes independent of ancestry, and many others may have distributional differences by ancestry but are equally valid after centering with respect to a readily collected population reference.

In contrast, PRS are uniformly less useful in understudied populations due to differences in genomic variation and population history^13,14^. No analogous solution of defining ethnicity-specific reference intervals would ameliorate health disparities implications for PRS or fundamentally aid interpretability in non-European populations. Rather, as we and others demonstrate, PRS are unique in that even with multi-ethnic population references, these scores are fundamentally less informative in populations more diverged from GWAS study cohorts.

The clinical use and deployment of genetic risk scores needs to be informed by the issues surrounding tests that currently would unequivocally provide much greater benefit to the subset of the world’s population which is already on the positive end of healthcare disparities. Conversely, African descent populations, which already endure many of the largest health disparities globally, are often predicted marginally better, if at all, compared to random (**Figure 4F**). They are therefore least likely to benefit from improvements in precision healthcare delivery from genetic risk scores with existing data due to human population history and study biases. This is a major concern globally and especially in the U.S., which already leads other middle- and high-income countries in both real and perceived healthcare disparities^69,70^. Thus, we would strongly urge that any discourse on clinical use of PRS include a careful, quantitative assessment of the economic and health disparities impacts on underrepresented populations that might be unintentionally introduced, and raise awareness about how to eliminate these disparities.

## How do we even the ledger?

What can be done? The single most important step towards parity in PRS accuracy is by vastly increasing the diversity of participants included and analyzed in genetic studies, which will improve utility for all and most rapidly for underrepresented groups. Regulatory protections against genetic discrimination are necessary to accompany calls for more diverse studies; while some already exist in the U.S., including for health insurance and employment opportunities via the Genetic Information Nondiscrimination Act (GINA), stronger protections in these and other areas globally will be particularly important for minorities and/or marginalized groups. An equal investment in GWAS across all major ancestries and global populations is the most obvious solution to generate a substrate for equally informative risk scores but is not likely to occur any time soon absent a dramatic priority shift given the current imbalance and stalled diversifying progress over the last five years (**Figure 1**, **Supplementary Figure 1**). While it may be challenging or in some cases infeasible to acquire sample sizes large enough for PRS to be equally informative in all populations, some much-needed efforts towards increasing diversity in genomics that support open sharing of GWAS summary data from multiple ancestries are underway. Examples include the *All of Us* Research Program, the Population Architecture using Genomics and Epidemiology (PAGE) Consortium, as well as some disease-focused consortia, such as the T2D-genes and Stanley Global initiatives on the genetics of type II diabetes and psychiatric disorders, respectively. Supporting data resources such as imputation panels, multi-ethnic genotyping arrays, gene expression datasets from genetically diverse individuals, and other tools are necessary to similarly empower these diverse studies for all populations. The lack of supporting resources for diverse ancestries creates financial challenges for association studies with limited resources, e.g. raising questions about whether to genotype samples on GWAS arrays that may favor European allele frequencies versus sequence samples, and how dense of an array to choose or how deeply to sequence^71,72^.

Additional leading global efforts also provide easy unified access linking genetic, clinical record, and national registry data in more homogeneous continental ancestries, such as the UK Biobank, BioBank Japan, China Kadoorie Biobank, and Nordic efforts (e.g. in Danish, Estonian, Finnish, and other integrated biobanks). Notably, some of these biobanks such as UK Biobank have participants with considerable global genetic diversity that enables multi-ethnic comparisons; although minorities from this cohort provide the largest deeply phenotyped GWAS cohorts for several ancestries, these individuals are often excluded in current statistical analyses in favor of single ancestries, large sample sizes, and the simplicity afforded by genetic homogeneity. These considerations notwithstanding, there are critical needs and challenges for expanding the scale of genetic studies of heritable traits in diverse populations; this is especially apparent in Africa where humans originated and retain the most genetic diversity, as Africans are understudied but disproportionately informative for genetic analyses and evolutionary history^27,73^. The most notable investment here comes from the Human Heredity and Health in Africa (H3Africa) Initiative, increasing genomics research capacity in Africa through more than $216 million in funding from the NIH (USA) and Wellcome Trust (UK) for genetics research led by African investigators^55,74^. The increasing interest and scale of genetic studies in low- and middle-income countries (LMICs) raises ethical and logistical considerations about data generation, access, sharing, security, and analysis, as well as clinical implementation to ensure these advances do not only benefit high-income countries. Frameworks such as the H3ABioNet, a pan-African bioinformatics network designed to build capacity to enable H3Africa researchers to analyze their data in Africa, provide cost-effective examples for training local scientists in LMICs^75^.

The prerequisite data for dramatically increasing diversity also hypothetically exist in several large-scale publicly funded datasets such as the Million Veterans Project and Trans-Omics for Precision Medicine (TOPMed), but with problematic data access issues in which even GWAS summary data within and across populations are not publicly shared. Existing GWAS consortia also need to carefully consider the granularity of summary statistics they release, as finer scale continental ancestries and phenotypes in large, multi-ethnic projects enable ancestry-matched analyses not possible with a single set of summary statistics. While there is an understandable patient privacy balance to strike when sharing individual-level data, GWAS summary statistics from all publicly funded and as many privately funded projects as possible should be made easily and publicly accessible to improve global health outcomes. Efforts to unify phenotype definitions, normalization approaches, and GWAS methods among studies will also improve comparability.

To enable progress towards parity, it will be critical that open data sharing standards be adopted for all ancestries and for genetic studies of all sample sizes, not just the largest European results. Locally appropriate and secure genetic data sharing techniques as well as equitable technology availability will need to be adopted widely in Asia and Africa as they are in Europe and North America, to ensure that maximum value is achieved from existing and ongoing efforts that are being developed to help counter the current imbalance. Simultaneously, ethical considerations require that research capacity is increased in LMICs with simultaneous growth of diverse population studies to balance the benefits of these studies to scientists and patients globally versus locally to ensure that everyone benefits. Methodological improvements that better define risk scores by accounting for population allele frequency, LD, and/or admixture differences appropriately are underway and may help considerably but will not by themselves bring equality. All of these efforts are important and should be prioritized not just for risk prediction but more generally to maximize the use and applicability of genetics to inform on the biology of disease. Given the acute recent attention on clinical use of PRS, we believe it is paramount to recognize their potential to improve health outcomes for all individuals and many complex diseases. Simultaneously, we as a field must address the disparity in utility in an ethically thoughtful and scientifically rigorous fashion, lest we inadvertently enable genetic technologies to contribute to, rather than reduce, existing health disparities.

## Supporting information

Supplementary Data Sets 1-3

## Author contributions

A.R.M. and M.J.D. conceived and designed the experiments. A.R.M. and M.K. performed statistical analysis. A.R.M. and M.K. analyzed the data. A.R.M., M.K., Y.K., Y.O., B.M.N., and M.J.D. contributed reagents/materials/analysis tools. A.R.M., M.K., B.M.N., and M.J.D. wrote the paper.

## Acknowledgments

We thank Amit Khera for helpful discussions. We also thank Michiaki Kubo, Yoshinori Murakami, Masato Akiyama, and Kazuyoshi Ishigaki for their support in the BioBank Japan Project analysis. We are grateful to Steven Gazal for his help in calculating LD scores. This work was supported by funding from the National Institutes of Health (K99MH117229 to A.R.M.). UK Biobank analyses were conducted via application 31063. The BioBank Japan Project was supported by the Tailor-Made Medical Treatment Program of the Ministry of Education, Culture, Sports, Science, and Technology (MEXT) and the Japan Agency for Medical Research and Development (AMED). M.K. was supported by a Nakajima Foundation Fellowship and the Masason Foundation.

