## Supplementary Note, Supplementary Tables 1-9, Supplementary Figures 1-13 for "Current clinical use of polygenic scores will risk exacerbating health disparities"

**Supplementary Material**

**Supplementary Note**

### 1. Methods

#### GWAS catalog analyses

*Ancestry over time*

We downloaded the GWAS catalog report named “All ancestry data” on 7/17/2018. We excluded individuals whose ancestry was listed as “Not reported.” Studies were sorted by date, and cumulative number of individuals over time were calculated. Individuals were grouped by broad ancestral categories, as defined previously[1].

*Frequency by ancestry*

We downloaded the GWAS catalog (v1.0.2) on 8/14/2018. To assess the frequency of GWAS catalog variants in African, East Asian, and European continental populations, we extracted African (AFR), East Asian (EAS), and European (EUR) individuals from the phase 3 1000 Genomes Project, where AFR excluded African Caribbeans in Barbados (ACB) and Americans of African Ancestry in SW USA (ASW) populations due to their recent European ancestry. We then computed minor allele frequency with plink2[2].

#### GWAS methods

*Quantitative traits*

GWAS for both the UK Biobank (UKBB) and BioBank Japan (BBJ) data were conducted using the same sample sizes for each phenotype (**Supplementary Table 6**). BBJ was ascertained on 47 diseases that likely influence or are correlated with some blood panel or anthropometric traits[3]. In BBJ, we thus first withheld a randomly selected 5,000 samples matching case/control fraction across diseases for prediction and used the rest of the cohort for conducting GWAS of the anthropometric and blood panel traits. Because the UKBB includes more individuals with homogeneous ancestry (N=361,195)[4], we matched sample sizes to the smaller BBJ data. All phenotypes evaluated were processed using PHESANT, as described previously[4], which curated and transformed phenotypes into normally distributed quantitative traits and categorical variables. Because basophil and eosinophil counts were binned with PHESANT resulting in lower observed $h^2$, we rank normalized these phenotypes separately. We ran all GWAS in Hail (https://hail.is/) using v0.1. As covariates, we included age, sex, $age^2$, age * sex, $age^2$ * sex, and the first 20 principal components. Because of the BBJ disease ascertainment strategy[3], we included additional indicator variables for each disease for BBJ only.

*Disease endpoints*

In addition to the 17 quantitative traits described above, we also examined five disease endpoints which exist in both BBJ and UKBB. In UKBB, phenotypes were defined based on PheCodes[5] using ICD-10 and ICD-9 primary and secondary diagnoses (UKBB Fields 41202–41204). In BBJ, the corresponding ICD codes were manually mapped to available medical records in their database (**Supplementary Data Set 1**). We conducted GWAS for both BBJ and UKBB using the same sample sizes for cases and controls (**Supplementary Table 8**). We first withheld a randomly selected 500 cases and 500 controls from each cohort, and then matched numbers of cases and controls. All GWAS were conducted using logistic regression in Hail v0.1 with the same covariates: age, sex, $age^2$, age * sex, $age^2$ * sex, and the first 20 principal components.

*Global structure in the UK Biobank*

To assess global population structure in the UK Biobank, we used the 1000 Genomes Project phase 3 data to assign "super population" labels defined in 1000 Genomes. Briefly, we intersected genotyped SNPs in the UK Biobank with SNPs genotyped in 1000 Genomes, filtering to SNPs with a minor allele frequency > 5%, excluding indels, removing ambiguous (i.e. A->T, T-> A, C->G, or G->C) SNPs, filtering to missingness < 1%, and pruning for LD $R^2$ < 0.2, resulting in 162,114 biallelic intersecting SNPs. Using this set of SNPs, we computed PCA for the 2,504 1000 Genomes individuals, then projected 488,377 UK Biobank individuals into the same PCA space. As in the ExAC project[6], we used a random forest classifier to assign super population labels based on the first 6 PCs (https://github.com/macarthur-lab/gnomad_hail/blob/master/utils/generic.py#L778). Counts of individuals by super population are shown in **Supplementary Table 7**.

*African ancestry individuals*

A more stringent set of filters was applied compared to the procedure described above for PRS analyses on African ancestry individuals in **Figure 4** to obtain a more ancestrally homogeneous set of individuals. These stricter filters were used for consistency with the Neale Lab UKBB GWAS of European ancestry individuals (**Supplementary Table 7**) and because considerable diversity in ancestral origins is present among Africans ancestry individuals in UKBB. We first performed PCA on an unrelated set of 1,919 individuals of African descent from the intersection between 1000 Genomes Project phase 3 Omni2.5 genotype data (AFR super population, N=627) and African Genome Variation Project Omni2.5 genotype data (N=1,292), computing the first 20 PCs using Hail 0.1. We projected all UK Biobank individuals assigned to the AFR super population in the "*Global structure in the UK Biobank*" section above onto this PC space. We then excluded related individuals. For these UKBB AFR samples, we then adapted the PCA-based European ancestry sample selection criteria for UK Biobank described here:
https://github.com/Nealelab/UK_Biobank_GWAS/blob/master/ukb31063_eur_selection.R.
Briefly, because of the greater ancestral heterogeneity, we used this sample selection to use code to draw ellipses in each PC space but narrowed the selection criteria to include individuals within two standard deviations along the first 10 PCs, keeping 5,739 of the 8,503 unrelated AFR individuals for PRS analyses in **Figure 4,** as shown in **Supplementary Figure 11**.

**Polygenic risk score (PRS) methods**

To ensure that both datasets started with the same number of SNPs, we extracted the intersecting SNP set across datasets (N=5,178,318 SNPs), then clumped the UKBB and BBJ summary statistics using EUR and EAS super populations from the phase 3 1000 Genomes Project data, respectively. Briefly, we used plink to clump variants using the following flags: --clump-p1 0.01 --clump-p2 1 --clump-r2 0.5 --clump-kb 250. We computed polygenic risk scores using Hail v0.2 for SNPs meeting several p-value thresholds: 5e-8, 1e-6, 1e-4, 1e-3, and 1e-2. We evaluated PRS accuracy in independent individuals from each biobank who were withheld from the GWAS. To assess genetic prediction accuracy, we computed partial $R^2$ attributable to the PRS from nested models, in which the full linear model was the true phenotype ~ PRS + all covariates described in *"GWAS methods"* above, and the nested model dropped only the PRS term. We bootstrapped individual phenotypes with their covariates to compute 95% confidence intervals on $R^2$ values. For disease endpoints, we instead used a logistic regression model and computed Nagelkerke's and liability-scale pseudo-$R^2$ as in Lee, SH., *et al*[7]. We set the population prevalence of each disease as 0.84% (atrial fibrillation), 0.0911% (colorectal cancer), 3.3% (glaucoma), 0.46% (rheumatoid arthritis), and 5.8% (type 2 diabetes) for both UKBB and BBJ based on the median of the reported values from the Incidence and Prevalence Database by Clarivate Analytics IPD (http://www.tdrdata.com). All computed PRS statistics including $R^2$ values (liability-scale and Nagelkerke's R2 for binary traits), 95% confidence intervals, and p-values are in **Supplementary Data Sets 2 and 3** for all traits, discovery and target populations, and GWAS p-value thresholds.

We note that although BBJ tended to show lower $R^2$ values than UKBB (**Figure 4, Supplementary Figures 2-3**) for quantitative traits (and vice versa for disease endpoints), we only consider these values for comparison within the same cohort, as they are not always easily comparable across different study cohorts. The observed lower $R^2$ in BBJ may be attributed to various factors, including lower observed heritability in BBJ than UKBB for these traits even when using the same sample sizes (**Supplementary Table 2**), baseline cohort characteristics (e.g., healthy volunteers for UKBB and diseases patients for BBJ), and numerous environmental factors (e.g. in the UK versus Japan).

*Relative prediction accuracy*
We compared prediction accuracy of the 17 traits in **Supplementary Table 6** across populations, as displayed in **Figure 3**. We computed $R^2$ values for each trait and for several p-value thresholds as described above for each of the genetically inferred continental-level populations (as described in "*Global structure in the UK Biobank*"). For each population and phenotype, we selected the predictor with the highest accuracy. Using these European-derived summary statistics, $R^2$ is highest for each trait in the holdout European individuals, as expected. We computed prediction accuracy relative to Europeans as $\frac{R^2_{population}}{R^2_{European}}$.

### Heritability estimates from summary statistics
We applied LD score regression[8] to all 17 quantitative traits and five diseases in UKBB and BBJ for which we generated summary statistics with matched sample sizes to estimate the heritability explained by the genome-wide high-quality common SNPs present in the HapMap 3 reference panel[9]. We estimated heritability using the default LD score[8] (without any functional annotations) and the baseline LD score (v2.1) which includes 86 functional annotations[10]. We used population-matched LD score references (i.e., EUR for UKBB and EAS for BBJ) downloaded from the authors' website (https://data.broadinstitute.org/alkesgroup/LDSCORE/). The major histocompatibility complex (MHC) region (chromosome 6: 25–34 Mb) was excluded from the analysis because of its complex LD structure.

### Trans-ethnic genetic correlation
Trans-ethnic genetic correlation compares the estimated correlation of common variant effect sizes at SNPs common in two populations. We computed trans-ethnic genetic correlation for

all 17 quantitative traits and five diseases between UKBB and BBJ using Popcorn[11]. Trans-ethnic genetic correlation was calculated for both genetic effect and genetic impact as defined previously[11]. For Hb, Ht, and Neutrophil, we computed trans-ethnic genetic correlation using regression ('--use_regression' option) rather than maximum likelihood (default), as the default approach produced unstable estimates for these (ρg=1, SE=0, and p=0). We used pre-computed cross-population scores for EUR and EAS 1000 Genomes populations provided by the authors (https://www.dropbox.com/sh/37n7drt7q4sjrzn/AAAa1HFeeRAE5M3YWG9Ac2Bta). For Popcorn analysis, we used 3,012,341 intersecting SNPs which exist in the pre-computed scores and both UKBB and BBJ's summary statistics and are not in the MHC region (chromosome 6: 25–34 Mb).

### 2. PRS accuracy as a function of age

Like other existing biomarkers, the predictive utility of PRS may change as a function of age, consistent with age-dependent heritability for some traits[12]. For example, increasing age is associated with higher risk of coronary artery disease, and higher PRS accelerate this increased risk[13]. Consequently, the age of intervention e.g. with statins needs to be evaluated in aggregate with other clinical risk factors that change over time. Autism spectrum disorder and schizophrenia also have a genetic basis with differing developmental trajectories; their shared genetic influences decrease with age, whereas the genetic overlap between schizophrenia and social communication difficulties persists with age[14]. Further work on how prediction accuracy varies as a function of age across phenotypes is needed.

### 3. Previous studies of PRS generalizability across populations

*PRS prediction using European GWAS summary statistics*

We have assembled prediction accuracy statistics from several studies using the largest European GWAS to predict several phenotypes in target European and non-European cohorts. For example, multiple schizophrenia studies consistently predicted risk on average 2.2-fold worse in East Asians relative to Europeans, (i.e. μ=0.46, σ=0.06), using summary statistics from a Eurocentric GWAS[15,16] (**Supplementary Figure 13**), despite the fact that there is no significant genetic heterogeneity in schizophrenia between the two populations[17]. This finding is even more pronounced in African Americans, consistent with higher genetic divergence from Europeans than between Europeans and East Asians[18]. Across several phenotypes with a range of genetic architectures in which empirical evaluations were available, including BMI, educational attainment, height, and schizophrenia, prediction accuracy using European GWAS summary statistics was on average 4.5-fold less accurate in African Americans than in Europeans (i.e. μ=0.22, σ=0.09, **Supplementary Figure 13**)[16,19-23]. By extension, prediction accuracy is expected to be even lower in African Americans with higher than average African ancestry or among populations with greater divergence from Europeans (e.g. some southern African populations).

*Promise from diverse GWAS to improve PRS accuracy across populations*

Several GWAS conducted outside European populations have disproportionately improved PRS accuracy in ancestry-matched individuals. These results suggest that diverse population GWAS are likely to improve PRS accuracy for all populations, with especially rapid improvement for underrepresented populations. For example, a BioBank Japan (BBJ) GWAS study (N=158,284) showed that compared to a 2× larger European GWAS (N=322,154), the variance in BMI explained in an independent Japanese cohort with Japanese GWAS summary statistics was on average 1.5-fold greater than with European GWAS summary statistics ($R^2$=0.154 vs 0.104 at $p < 0.05$, respectively)[24]. Similarly, a Chinese schizophrenia study (N=7,699 cases and 18,327 controls) showed that compared to an effectively 5.1× larger European GWAS (N=36,989 cases, 113,075 controls), prediction accuracy in an independent Chinese cohort with GWAS summary statistics from China far surpassed prediction accuracy from European summary statistics by 2.63 fold (2.34% versus 6.16%)[25]. Relatedly, an East Asian schizophrenia study (N=22,778 cases and 35,362 controls) showed that compared to an effectively 3× larger European study, prediction accuracy in East Asians was on average 1.3-fold higher than with European summary statistics (liability $R^2$=0.029 vs 0.022, respectively)[17].

#### 4. Effect sizes estimates differ, even if causal effects are the same

The marginal GWAS estimate differs because LD varies across populations. Mathematically, this is defined as:

$$\hat{\beta}_j = \sum_{k=1}^{m} r_{j,k}\beta_k + \epsilon_j$$

where $\hat{\beta}_j$ are effect size estimates at SNP $j$, $r_{j,k}$ is pairwise SNP LD between SNPs $j$ and $k$, $\beta_k$ is the causal SNP effect at nearby SNP $k$, and $\epsilon$ is residual error from bias or noise. More simply, when causal effects are the same across populations, effect size estimates at SNPs tagging these causal variants from which we construct predictors will differ across populations.

#### 5. Genetically determined ancestry versus self-identified race/ethnicity

For diverse genetic studies in the U.S. and globally, genetically determined ancestry and populations are important to delineate from self-identified race/ethnicity. Only controlling for the former can account for stratification of allele frequencies within a population. The latter may provide additional information about environmental racial correlates.

#### 6. Considerations of uneven population sample sizes and PRS accuracy

To maximally benefit all populations, the largest existing GWAS results should be used. Down-sampling the largest European GWAS for the sake of parity results in worse predictors for all individuals.

#### 7. Software availability

Code used to generate the results in this manuscript can be accessed here: https://github.com/armartin/prs_disparities

#### 8. Data availability

All data used in this study were available through previous work. As described in the Life Sciences Reporting Summary, UK Biobank analyses were conducted via application 31063. BBJ GWAS summary statistics are publicly available at our website (http://jenger.riken.jp/en/) and the National Bioscience Database Center (NBDC) Human Database (Research ID: hum0014). Genotype data from the BBJ subjects was deposited at the NBDC Human Database (Research ID: hum0014).

**Supplementary Table 1. $R^2$ measures across populations from European GWAS.** For binary traits, liability-scale $R^2$ is reported where possible and Nagelkerke's $R^2$ is reported elsewhere.

| Study population | Target population | Target cohort | Phenotype | $R^2$ | Relative to European | Reference |
|---|---|---|---|---|---|---|
| European | European | HRS | BMI | 0.058 | N/A | Ware et al, 2017 |
| European | African American | HRS | BMI | 0.015 | 0.26 | Ware et al, 2017 |
| European | European | ARIC | BMI | 0.016 | N/A | Belsky et al, 2013 |
| European | African American | ARIC | BMI | 0.001 | 0.09 | Belsky et al, 2013 |
| European | European | Add Health | EA | 0.032 | N/A | Domingue et al, 2015 |
| European | African American | Add Health | EA | 0.012 | 0.37 | Domingue et al, 2015 |
| European | European | HRS | EA | 0.060 | N/A | Ware et al, 2017 |
| European | African American | HRS | EA | 0.010 | 0.17 | Ware et al, 2017 |
| European | European | HRS | EA | 0.106 | N/A | Lee et al, 2018 |
| European | African American | HRS | EA | 0.016 | 0.15 | Lee et al, 2018 |
| European | European | HRS | Height | 0.104 | N/A | Ware et al, 2017 |
| European | African American | HRS | Height | 0.025 | 0.24 | Ware et al, 2017 |
| European | European | Multiple | SCZ | 0.085 | N/A | Ripke et al, 2014 |
| European | East Asian | JPN1 | SCZ | 0.046 | 0.55 | Ripke et al, 2014 |
| European | East Asian | TCR1 | SCZ | 0.033 | 0.39 | Ripke et al, 2014 |
| European | East Asian | HOK2 | SCZ | 0.040 | 0.47 | Ripke et al, 2014 |
| European | European | MGS | SCZ | 0.069 | N/A | Vilhjalmsson et al, 2015 |
| European | East Asian | JPN1 | SCZ | 0.032 | 0.46 | Vilhjalmsson et al, 2015 |
| European | East Asian | TCR1 | SCZ | 0.034 | 0.48 | Vilhjalmsson et al, 2015 |
| European | East Asian | HOK2 | SCZ | 0.027 | 0.39 | Vilhjalmsson et al, 2015 |
| European | African American | AFAM | SCZ | 0.015 | 0.22 | Vilhjalmsson et al, 2015 |
| European | European | Multiple | SCZ | 0.093 | N/A | Vassos et al, 2017 |
| European | African American | Multiple | SCZ | 0.027 | 0.29 | Vassos et al, 2017 |

**Supplementary Table 2. Observed trait heritability of 17 quantitative traits in each cohort using LD score regression.** Abbreviations are the same as in **Supplementary Table 6**.

| **Trait** | **BBJ** | | | | **UKBB** | | | |
|---|---|---|---|---|---|---|---|---|
| | **Default LDSC** | | **S-LDSC (baselineLD v2.1)** | | **Default LDSC** | | **S-LDSC (baselineLD v2.1)** | |
| | **Observed $h^2$** | **SE** | **Observed $h^2$** | **SE** | **Observed $h^2$** | **SE** | **Observed $h^2$** | **SE** |
| Basophil | 0.0441 | 0.0121 | 0.0678 | 0.0110 | 0.0213 | 0.0050 | 0.0251 | 0.0098 |
| BMI | 0.1361 | 0.0087 | 0.1819 | 0.0114 | 0.1955 | 0.0090 | 0.2536 | 0.0101 |
| DBP | 0.0430 | 0.0051 | 0.0529 | 0.0081 | 0.0984 | 0.0068 | 0.1465 | 0.0090 |
| Eosinophil | 0.0586 | 0.0093 | 0.0709 | 0.0140 | 0.1354 | 0.0167 | 0.2113 | 0.0152 |
| Hb | 0.0452 | 0.0053 | 0.0654 | 0.0080 | 0.1054 | 0.0107 | 0.1704 | 0.0106 |
| Height | 0.3059 | 0.0187 | 0.4167 | 0.0203 | 0.3675 | 0.0208 | 0.4765 | 0.0211 |
| Ht | 0.0457 | 0.0056 | 0.0716 | 0.0083 | 0.0942 | 0.0093 | 0.1584 | 0.0105 |
| Lymphocyte | 0.0516 | 0.0073 | 0.0827 | 0.0119 | 0.1318 | 0.0118 | 0.2032 | 0.0139 |
| MCH | 0.1309 | 0.0184 | 0.1354 | 0.0146 | 0.1942 | 0.0210 | 0.1870 | 0.0134 |
| MCHC | 0.0481 | 0.0080 | 0.0747 | 0.0114 | 0.0402 | 0.0052 | 0.0577 | 0.0073 |
| MCV | 0.1447 | 0.0178 | 0.1601 | 0.0154 | 0.1994 | 0.0201 | 0.2107 | 0.0136 |
| Monocyte | 0.0448 | 0.0090 | 0.0776 | 0.0140 | 0.1331 | 0.0177 | 0.1982 | 0.0190 |
| Neutrophil | 0.0758 | 0.0097 | 0.1031 | 0.0140 | 0.1153 | 0.0131 | 0.1607 | 0.0132 |
| Platelet | 0.1260 | 0.0148 | 0.1818 | 0.0164 | 0.2012 | 0.0179 | 0.2481 | 0.0139 |
| RBC | 0.0818 | 0.0093 | 0.1101 | 0.0106 | 0.1586 | 0.0141 | 0.2119 | 0.0121 |
| SBP | 0.0574 | 0.0063 | 0.0761 | 0.0095 | 0.1041 | 0.0070 | 0.1531 | 0.0096 |
| WBC | 0.0778 | 0.0074 | 0.1078 | 0.0092 | 0.1286 | 0.0114 | 0.1994 | 0.0098 |

**Supplementary Table 3. Observed trait heritability of five diseases in each cohort using LD score regression.** Abbreviations are the same as in **Supplementary Table 8**.

| Trait | BBJ | | | | UKBB | | | |
|---|---|---|---|---|---|---|---|---|
| | Default LDSC | | S-LDSC (baselineLD v2.1) | | Default LDSC | | S-LDSC (baselineLD v2.1) | |
| | Observed $h^2$ | SE | Observed $h^2$ | SE | Observed $h^2$ | SE | Observed $h^2$ | SE |
| AFib | 0.0354 | 0.0098 | 0.0413 | 0.0062 | 0.0228 | 0.0039 | 0.0259 | 0.0055 |
| CRC | 0.0056 | 0.0027 | 0.0095 | 0.0055 | 0.0040 | 0.0024 | 0.0087 | 0.0053 |
| Glaucoma | 0.0075 | 0.0025 | 0.0212 | 0.0050 | 0.0121 | 0.0026 | 0.0180 | 0.0047 |
| RA | 0.0086 | 0.0027 | 0.0212 | 0.0050 | 0.0076 | 0.0023 | 0.0147 | 0.0044 |
| T2D | 0.0473 | 0.0050 | 0.0777 | 0.0077 | 0.0641 | 0.0046 | 0.1012 | 0.0073 |

**Supplementary Table 4. Liability-scale trait heritability of five diseases in each cohort using LD score regression.** Abbreviations are the same as in **Supplementary Table 8**.

| Trait | BBJ | | | | UKBB | | | |
|---|---|---|---|---|---|---|---|---|
| | Default LDSC | | S-LDSC (baselineLD v2.1) | | Default LDSC | | S-LDSC (baselineLD v2.1) | |
| | Liability-scale $h^2$ | SE | Liability-scale $h^2$ | SE | Liability-scale $h^2$ | SE | Liability-scale $h^2$ | SE |
| AFib | 0.1036 | 0.0288 | 0.1210 | 0.0183 | 0.0668 | 0.0115 | 0.0758 | 0.0161 |
| CRC | 0.0226 | 0.0108 | 0.0383 | 0.0222 | 0.0163 | 0.0097 | 0.0349 | 0.0214 |
| Glaucoma | 0.0711 | 0.0234 | 0.2009 | 0.0471 | 0.1153 | 0.0248 | 0.1710 | 0.0447 |
| RA | 0.0521 | 0.0163 | 0.1290 | 0.0304 | 0.0463 | 0.0139 | 0.0893 | 0.0270 |
| T2D | 0.1120 | 0.0119 | 0.1840 | 0.0182 | 0.1517 | 0.0108 | 0.2396 | 0.0173 |

**Supplementary Table 5. Trans-ethnic genetic correlation between BBJ and UKBB using Popcorn.** $\rho_{ge}$: trans-ethnic genetic effect correlation and $\rho_{gi}$: trans-ethnic genetic impact correlation as defined previously[11]. Abbreviations are the same as in **Supplementary Tables 6 and 8**. *For these three traits, we generated Popcorn estimates using regression rather than maximum likelihood (default), as the default approach produced unstable estimates for these ($\rho_g$=1, SE=0, and p=0).

| **Trait** | $\rho_{ge}$ | **SE** | **P** | $\rho_{gi}$ | **SE** | **P** |
|---|---|---|---|---|---|---|
| Basophil | 0.5945 | 0.1221 | 0.0009 | 0.6409 | 0.1370 | 0.0088 |
| BMI | 0.7474 | 0.0230 | 0.0 | 0.7237 | 0.0232 | 0.0 |
| DBP | 0.8354 | 0.0509 | 0.0012 | 0.8100 | 0.0508 | 0.0002 |
| Eosinophil | 0.9656 | 0.0707 | 0.6266 | 0.9483 | 0.0732 | 0.4800 |
| Hb* | 0.9741 | 0.0878 | 0.7682 | 0.9449 | 0.0935 | 0.5561 |
| Height | 0.6932 | 0.0172 | 0.0 | 0.6737 | 0.0172 | 0.0 |
| Ht* | 0.9012 | 0.0789 | 0.2102 | 0.8924 | 0.0890 | 0.2264 |
| Lymphocyte | 0.9777 | 0.0666 | 0.7380 | 0.9753 | 0.0747 | 0.7415 |
| MCH | 0.9727 | 0.0547 | 0.6175 | 0.9555 | 0.0660 | 0.5001 |
| MCHC | 0.9167 | 0.0910 | 0.3596 | 0.9195 | 0.1058 | 0.4469 |
| MCV | 0.9565 | 0.0487 | 0.3722 | 0.9409 | 0.0572 | 0.3013 |
| Monocyte | 0.9946 | 0.0788 | 0.9453 | 1.0000 | 0.0269 | 0.9999 |
| Neutrophil* | 0.9328 | 0.0707 | 0.3417 | 0.9404 | 0.0722 | 0.4085 |
| Platelet | 0.9068 | 0.0548 | 0.0891 | 0.8856 | 0.0532 | 0.0316 |
| RBC | 0.9819 | 0.0475 | 0.7039 | 0.9759 | 0.0532 | 0.6505 |
| SBP | 0.8469 | 0.0430 | 0.0004 | 0.8323 | 0.0445 | 0.0002 |
| WBC | 0.8922 | 0.0402 | 0.0074 | 0.8941 | 0.0422 | 0.0120 |
| AFib | 1.0000 | 0.0254 | 0.9999 | 0.9799 | 0.0955 | 0.8337 |
| CRC | 0.7410 | 0.2771 | 0.3498 | 0.7741 | 0.2992 | 0.4503 |
| Glaucoma | 0.8825 | 0.1826 | 0.5200 | 0.9149 | 0.2023 | 0.6740 |
| RA | 0.7984 | 0.2031 | 0.3209 | 0.8024 | 0.2136 | 0.3550 |
| T2D | 1.0000 | 0.0005 | 0.9931 | 1.0000 | 0.0309 | 0.9999 |

**Supplementary Table 6. Number of total individuals overall, in BBJ and UKBB GWAS, and in the holdout target datasets for 17 quantitative traits.** Clumps are independent loci with $p < 0.01$, which were computed using the plink, as described in the PRS methods above. Abbreviations are as follows: BMI = body mass index, DBP = diastolic blood pressure, Hb = hemoglobin, Ht = Hematocrit, MCH = mean corpuscular hemoglobin, MCHC = mean corpuscular hemoglobin concentration, MCV = mean corpuscular volume, RBC = red blood cell count, SBP = systolic blood pressure, WBC = white blood cell count. $N_{target}$ describes the number of individuals used for cross-biobank PRS analyses in **Figure 4**.

| Trait | $N_{total}$ (BBJ) | $N_{GWAS}$ (BBJ & UKBB) | $N_{target}$ (BBJ & UKBB) | UKBB code | # BBJ clumps | # UKBB clumps |
|---|---|---|---|---|---|---|
| Basophil | 87665 | 82665 | 5000 | 30160 | 8939 | 8690 |
| BMI | 155426 | 150426 | 5000 | 21001 | 19114 | 21339 |
| DBP | 137991 | 132991 | 5000 | 4079 | 9865 | 14213 |
| Eosinophil | 88675 | 83675 | 5000 | 30150 | 9266 | 13061 |
| Hb | 144653 | 139653 | 5000 | 30020 | 10483 | 16184 |
| Height | 156569 | 151569 | 5000 | 50 | 37216 | 31854 |
| Ht | 144947 | 139947 | 5000 | 30030 | 10554 | 15408 |
| Lymphocyte | 91157 | 86157 | 5000 | 30120 | 9400 | 13648 |
| MCH | 121249 | 116249 | 5000 | 30050 | 12598 | 15222 |
| MCHC | 128232 | 123232 | 5000 | 30060 | 10272 | 10074 |
| MCV | 122912 | 117912 | 5000 | 30040 | 13169 | 16354 |
| Monocyte | 90593 | 85593 | 5000 | 30130 | 10886 | 13452 |
| Neutrophil | 79287 | 74287 | 5000 | 30140 | 9150 | 12211 |
| Platelet | 140610 | 135610 | 5000 | 30080 | 14843 | 19259 |
| RBC | 145426 | 140426 | 5000 | 30010 | 12467 | 18069 |
| SBP | 137981 | 132981 | 5000 | 4080 | 11231 | 14562 |
| WBC | 146158 | 141158 | 5000 | 30000 | 12664 | 17581 |

**Supplementary Table 7. Numbers of individuals with a given ancestry in UK Biobank.** Target individuals describes the numbers of individuals used in **Figure 3**.

| Super population | Total | Unrelated | Target individuals | Neale Lab GWAS |
|---|---|---|---|---|
| EUR | 447206 | 370407 | 5000 | 350326 |
| SAS | 9950 | 9015 | 9015 | 0 |
| AFR | 9288 | 8503 | 8503 | 0 |
| AMR | 4724 | 4329 | 4329 | 1 |
| EAS | 2421 | 2306 | 2306 | 0 |
| other | 14788 | 12252 | N/A | 10867 |
| TOTAL | 488377 | 406812 | 29153 | 361194 |

**Supplementary Table 8. Number of total cases and controls overall, in BBJ and UKBB GWAS, and in the holdout target datasets for five diseases.** Clumps are independent loci with p < 0.01, which were computed using the plink, as described in the PRS methods above. Abbreviations are as follows: AFib = atrial fibrillation, CRC = colorectal cancer, RA = rheumatoid arthritis, T2D = type 2 diabetes.

| Trait | $N_{cases}$ (BBJ) | $N_{controls}$ (BBJ) | $N_{cases}$ (UKBB) | $N_{controls}$ (UKBB) | $N_{GWAS, cases}$ (BBJ & UKBB) | $N_{GWAS, controls}$ (BBJ & UKBB) | $N_{target, cases}$ (BBJ & UKBB) | $N_{target, controls}$ (BBJ & UKBB) | # BBJ clumps | # UKBB clumps |
|---|---|---|---|---|---|---|---|---|---|---|
| AFib | 8174 | 154407 | 12975 | 336589 | 7674 | 153907 | 500 | 500 | 8357 | 8957 |
| CRC | 6691 | 154121 | 3961 | 338191 | 3461 | 153621 | 500 | 500 | 6795 | 8048 |
| Glaucoma | 5122 | 164788 | 3883 | 351405 | 3383 | 164288 | 500 | 500 | 7304 | 8206 |
| RA | 4024 | 165886 | 3797 | 323214 | 3297 | 165386 | 500 | 500 | 8014 | 8222 |
| T2D | 36802 | 131404 | 15803 | 343663 | 15303 | 130904 | 500 | 500 | 10097 | 12293 |

**Supplementary Table 9. Numbers of thalassemia and sickle cell ICD-10 codes reported across all UKBB individuals**

| ICD-10 code | Number of individuals |
|---|---|
| *D56 Thalassaemia* | |
| D56.1 Beta thalassaemia | 8 |
| D56.3 Thalassaemia trait | 3 |
| D56.9 Thalassaemia, unspecified | 2 |
| *D57 Sickle-cell disorders* | |
| D57.0 Sickle-cell anaemia with crisis | 24 |
| D57.1 Sickle-cell anaemia without crisis | 34 |
| D57.2 Double heterozygous sickling disorders | 3 |
| D57.3 Sickle-cell trait | 1 |
| D57.8 Other sickle-cell disorders | 1 |

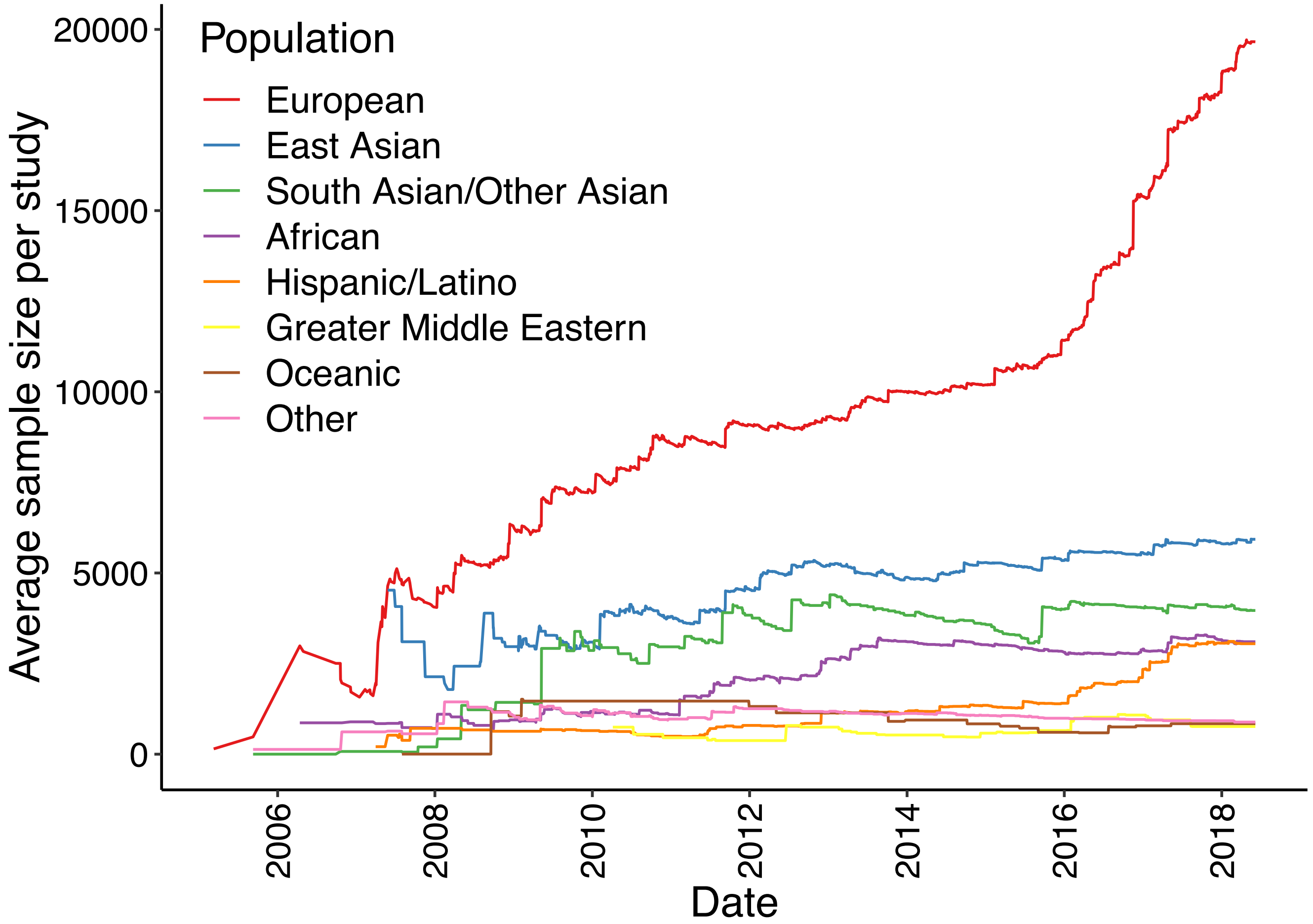


**Supplementary Figure 1. Average sample size per GWAS over time by population using GWAS catalog data.**

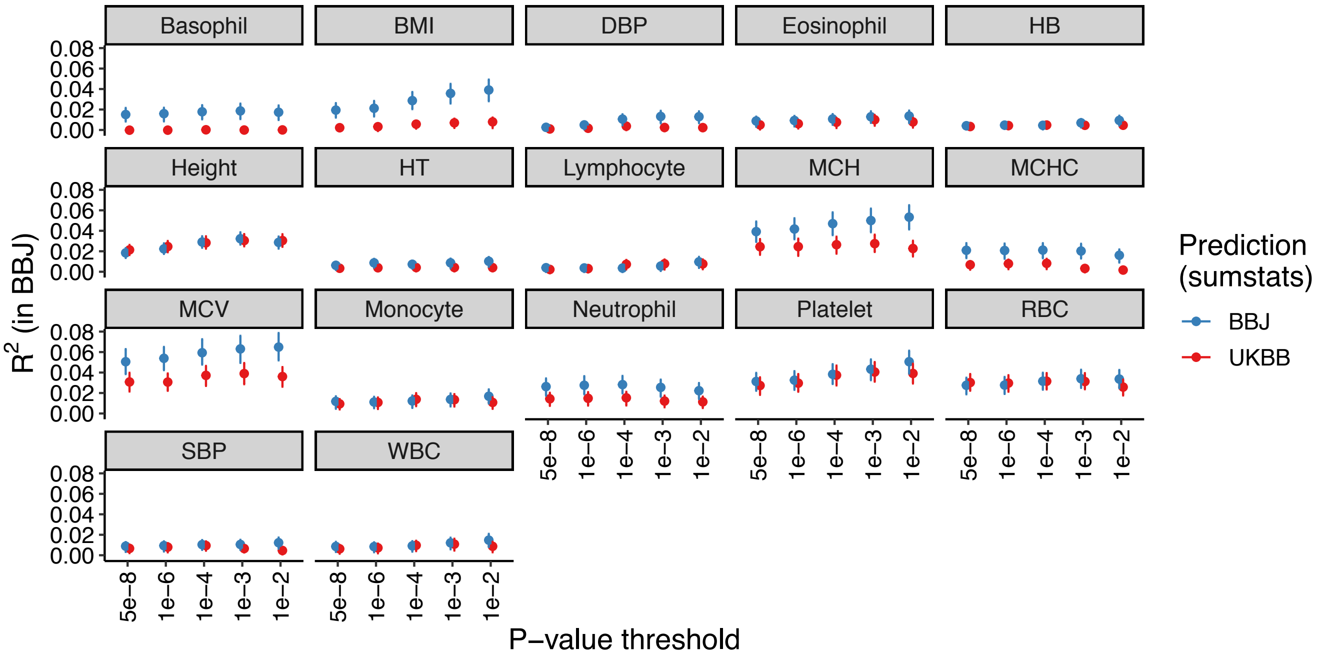


**Supplementary Figure 2. PRS accuracy for 17 quantitative traits under several p-value thresholds in BBJ.** Abbreviations are as in **Supplementary Table 6**. Points indicate the $R^2$ for each p-value threshold and lines correspond to 95% confidence intervals computed via bootstrap.

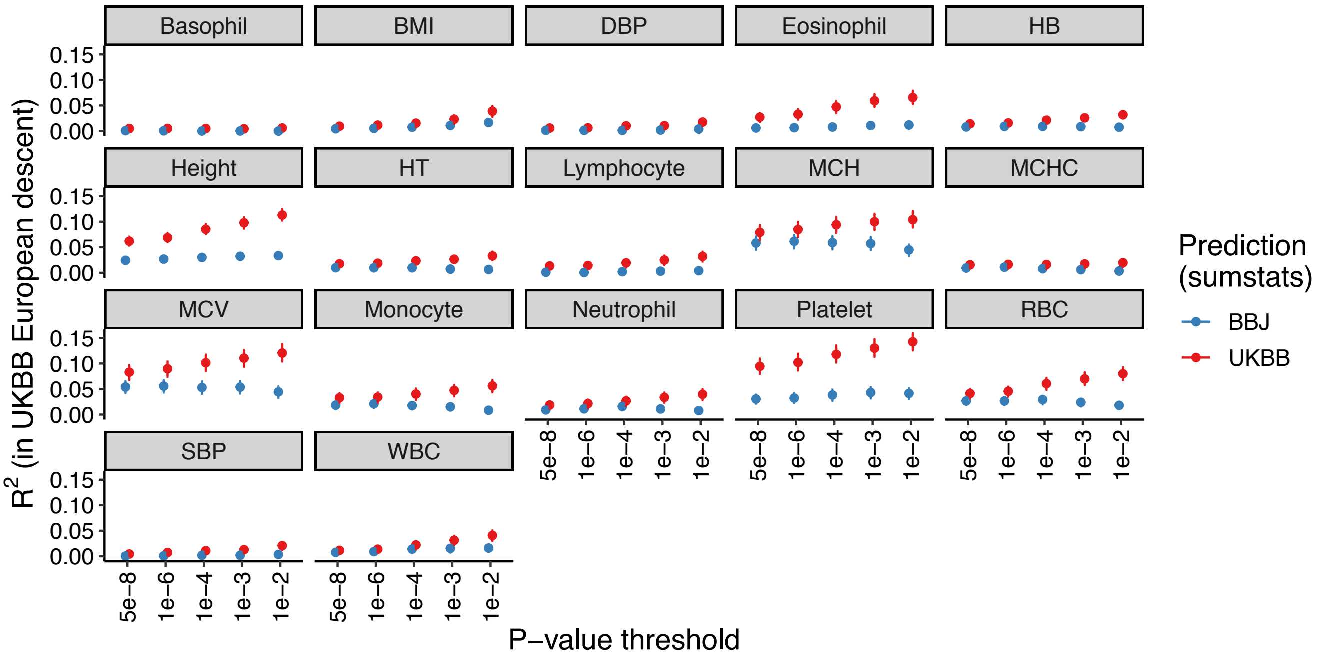


**Supplementary Figure 3. PRS accuracy for 17 quantitative traits under several p-value thresholds in UKBB individuals of European descent.** Abbreviations are as in **Supplementary Table 6**. Points indicate the $R^2$ for each p-value threshold and lines correspond to 95% confidence intervals computed via bootstrap.

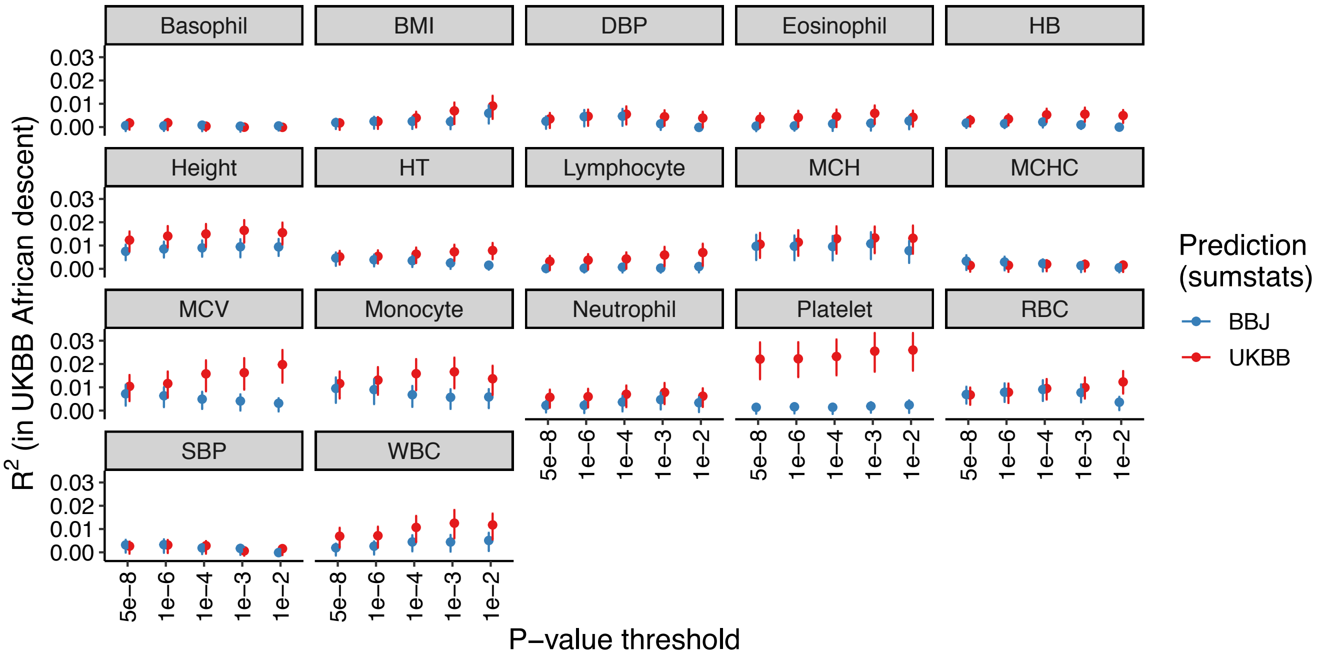


**Supplementary Figure 4. PRS accuracy for 17 quantitative traits under several p-value thresholds in UKBB individuals of African descent.** Abbreviations are as in **Supplementary Table 6**. Points indicate the $R^2$ for each p-value threshold and lines correspond to 95% confidence intervals computed via bootstrap.

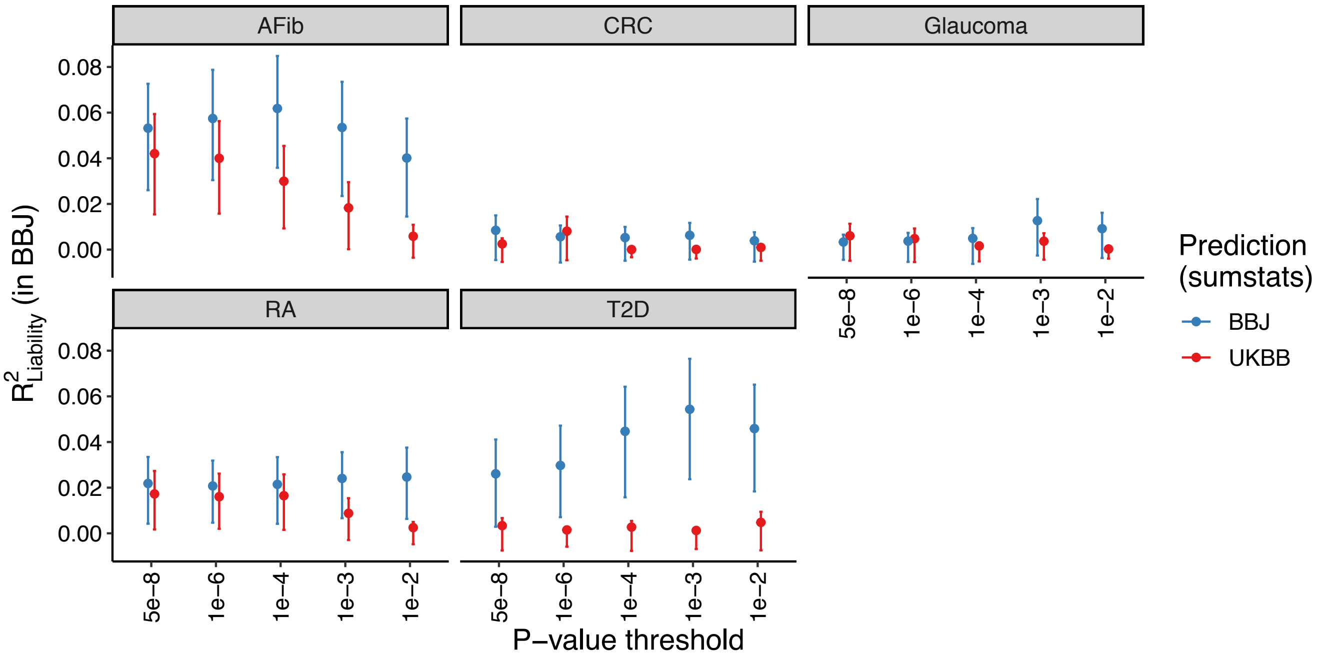


**Supplementary Figure 5. PRS accuracy for five diseases under several p-value thresholds in BBJ.** Abbreviations are as in **Supplementary Table 8**. Points indicate the liability $R^2$ for each p-value threshold and lines correspond to 95% confidence intervals computed via bootstrap.

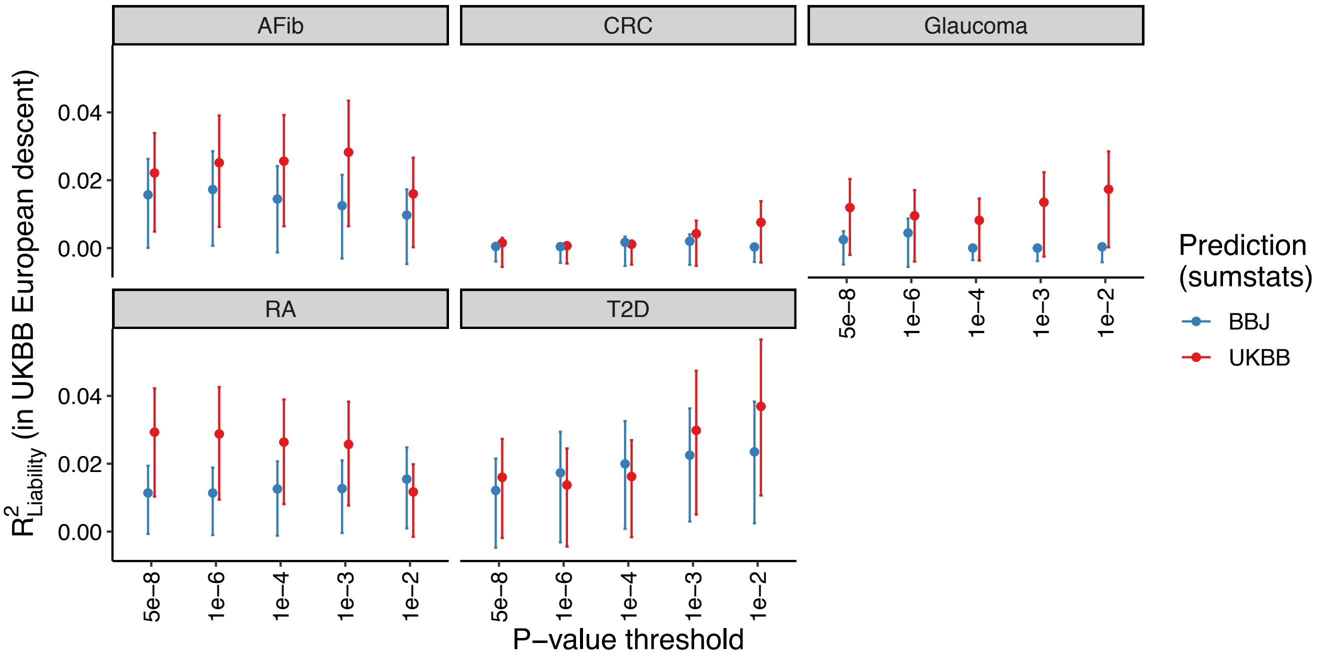


**Supplementary Figure 6. PRS accuracy for five diseases under several p-value thresholds in UKBB individuals of European descent.** Abbreviations are as in **Supplementary Table 8**. Points indicate the liability $R^2$ for each p-value threshold and lines correspond to 95% confidence intervals computed via bootstrap.

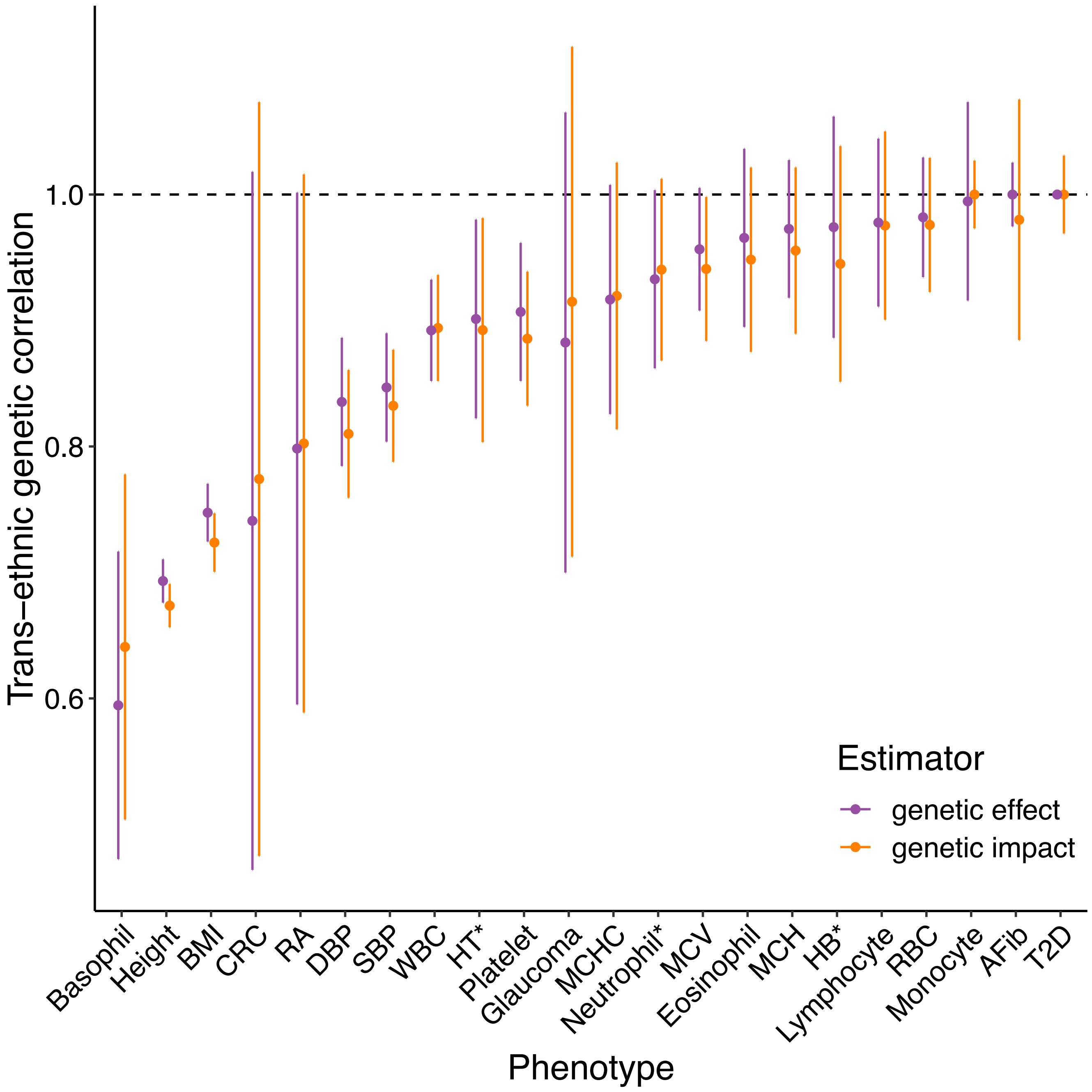


**Supplementary Figure 7. Trans-ethnic genetic correlation among BBJ and UKBB samples using Popcorn.** Abbreviations are as in **Supplementary Tables 6 and 8**. Results are as in **Supplementary Table 5**. Error bars indicate standard errors. *For these three traits, we generated Popcorn estimates using regression rather than maximum likelihood (default), as the default approach produced unstable estimates for these ($\rho_g$=1, SE=0, and p=0).

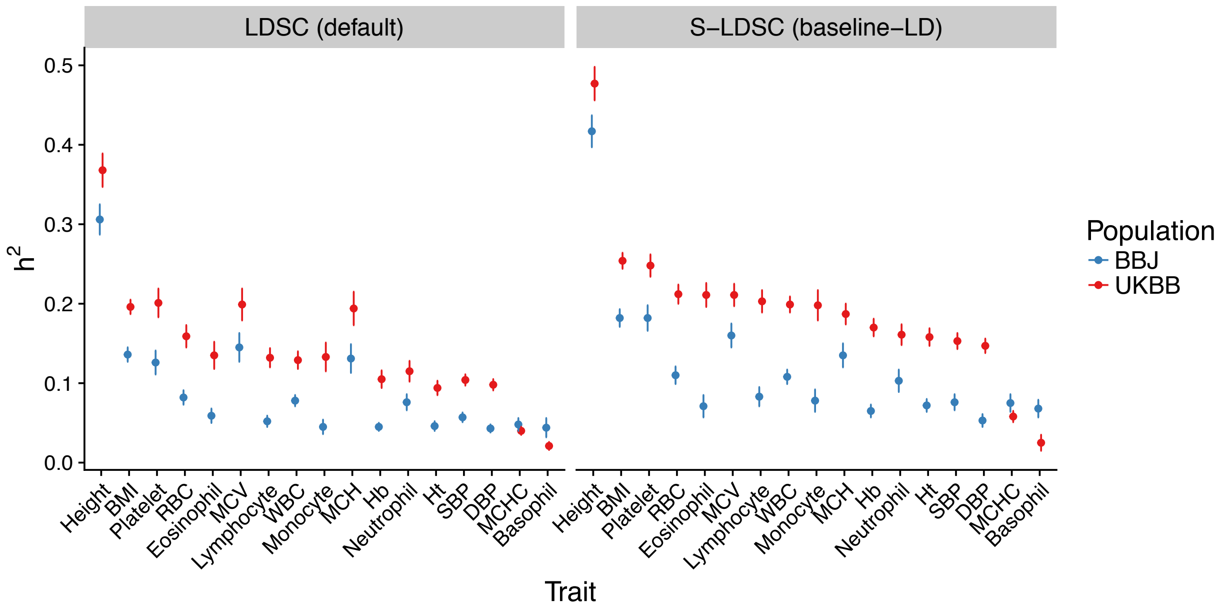


**Supplementary Figure 8. Estimated observed heritability in BBJ and UKBB across 17 quantitative traits using two LD score regression approaches.** The first approach (left) is the default for LD score regression, which doesn't include any stratification by functional categories, whereas the second approach (right) is the stratified LD score regression approach with the baseline LD model (v2.1) for each population. Abbreviations are as in **Supplementary Table 6**. Error bars indicate standard errors.

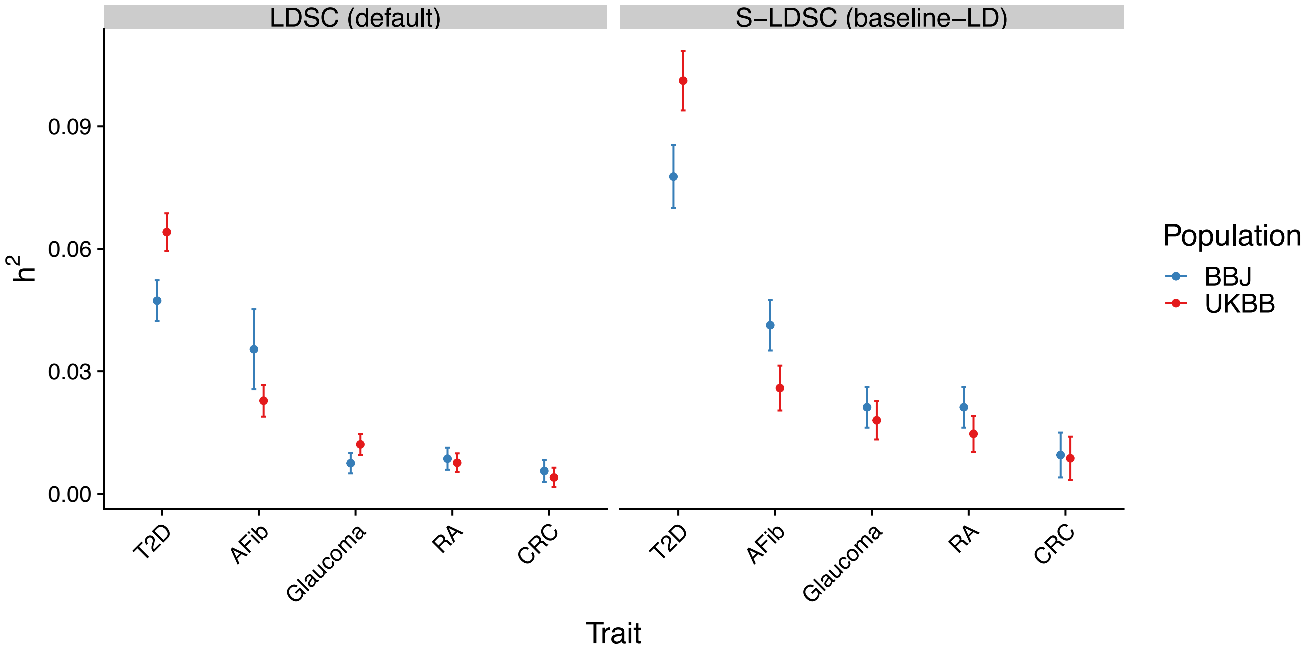


**Supplementary Figure 9. Estimated observed heritability in BBJ and UKBB across five diseases using two LD score regression approaches.** Abbreviations are as in **Supplementary Table 8**. Error bars indicate standard errors.

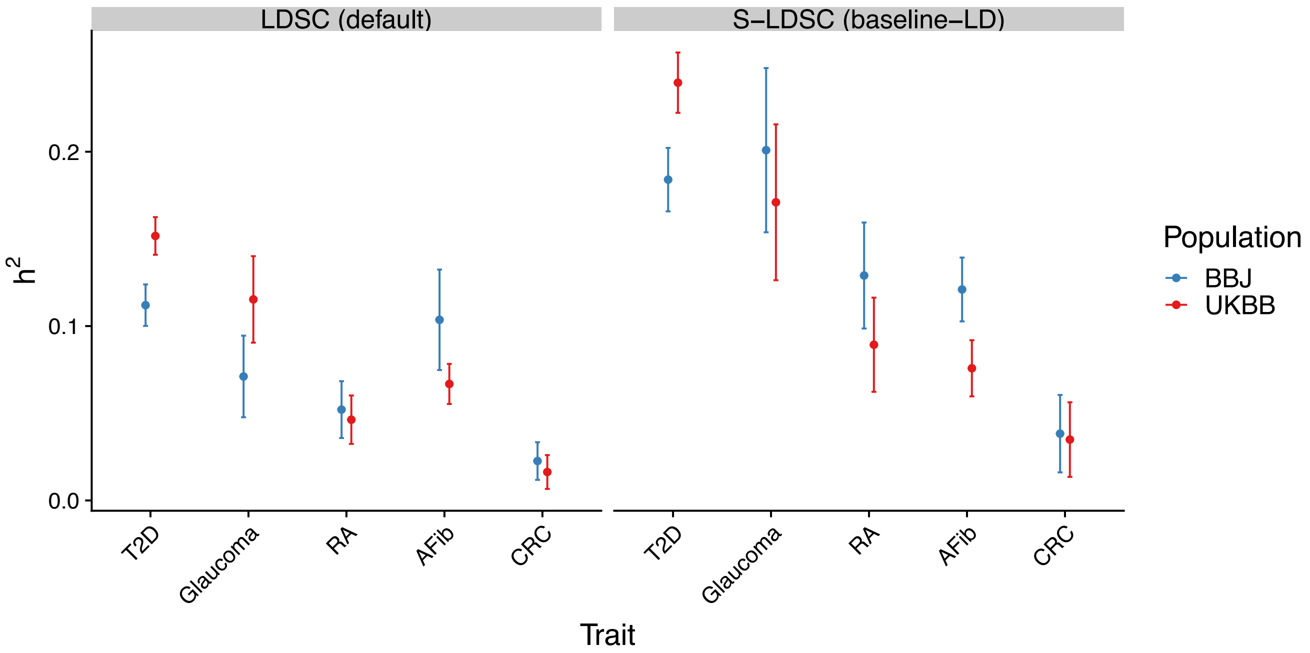


**Supplementary Figure 10. Estimated liability-scale heritability in BBJ and UKBB across five diseases using two LD score regression approaches.** Abbreviations are as in **Supplementary Table 8**. Error bars indicate standard errors.

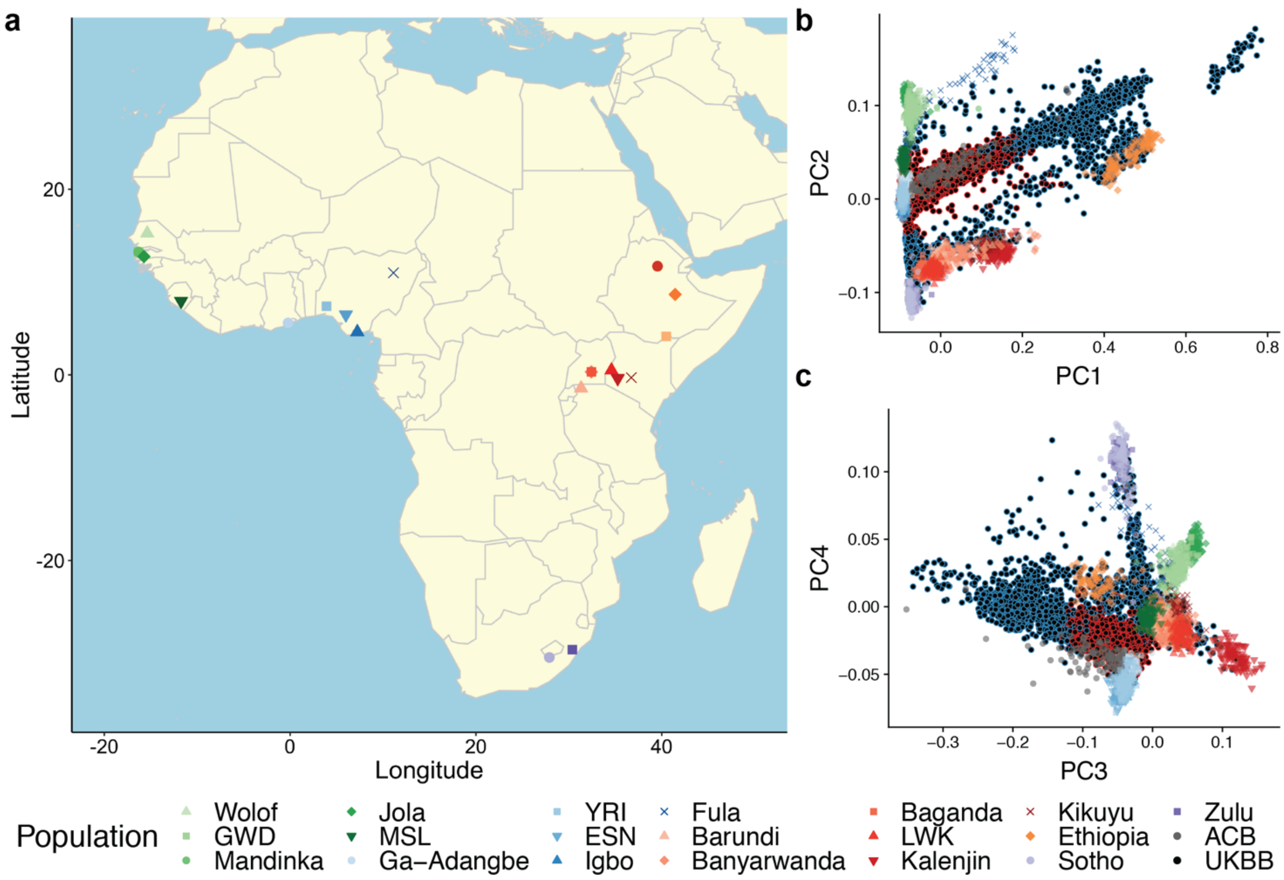


**Supplementary Figure 11. PCA of African descent UK Biobank individuals used for prediction accuracy assessment in Figure 4. a**) Map of Africa and approximate locations of reference panel individuals, including 1000 Genomes AFR populations (excluding ASW) and African Genome Variation Project (AGVP) populations. This map was created using open source data with R computing code. Three-letter abbreviations correspond to 1000 Genomes populations, as follows: GWD=Gambian in Western Divisions in the Gambia; MSL=Mende in Sierra Leone; YRI=Yoruba in Ibadan, Nigeria; ESN=Esan in Nigeria; LWK=Luhya in Webuye, Kenya; and ACB=African Caribbeans in Barbados. **b-c**) Reference panel individuals are plotted on top of UK Biobank Africans. The latter are shown in black circles, with a red outline if they were included in the PRS target samples in **Figure 4** and a blue outline if they were excluded. Plots show PCA using 1000 Genomes AFR + AGVP as reference individuals, and projecting UKBB African individuals into this PCA space for PC1 vs PC2 (**b**) and PC3 vs PC4 (**c**).

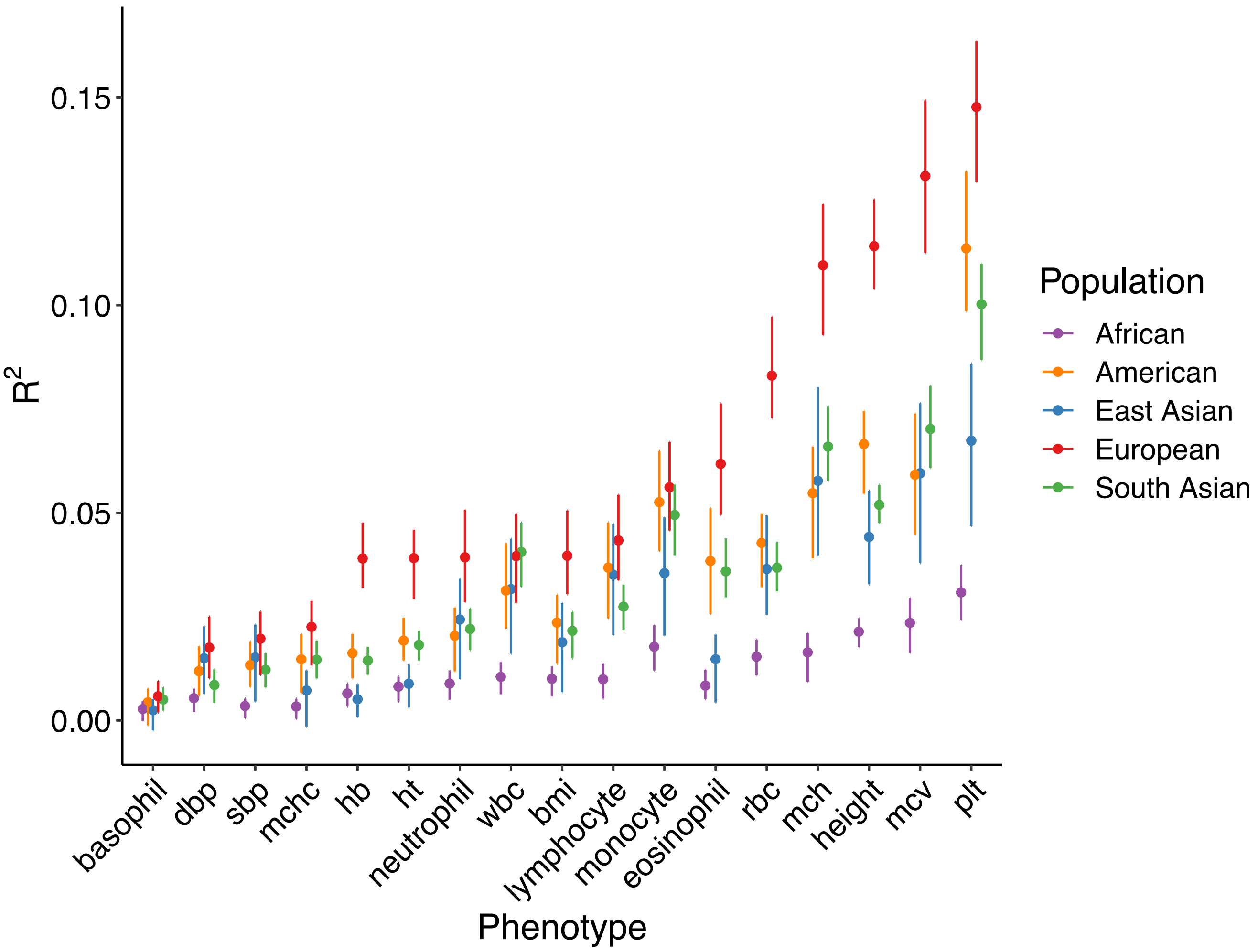


**Supplementary Figure 12. Prediction accuracy in UKBB in different populations for 17 phenotypes.** The best predictors among 5 p-value thresholds (**Supplementary Note**) are shown for each population. Abbreviations are as in **Supplementary Table 6**. Points indicate the $R^2$ for each p-value threshold and lines correspond to 95% confidence intervals computed via bootstrap.

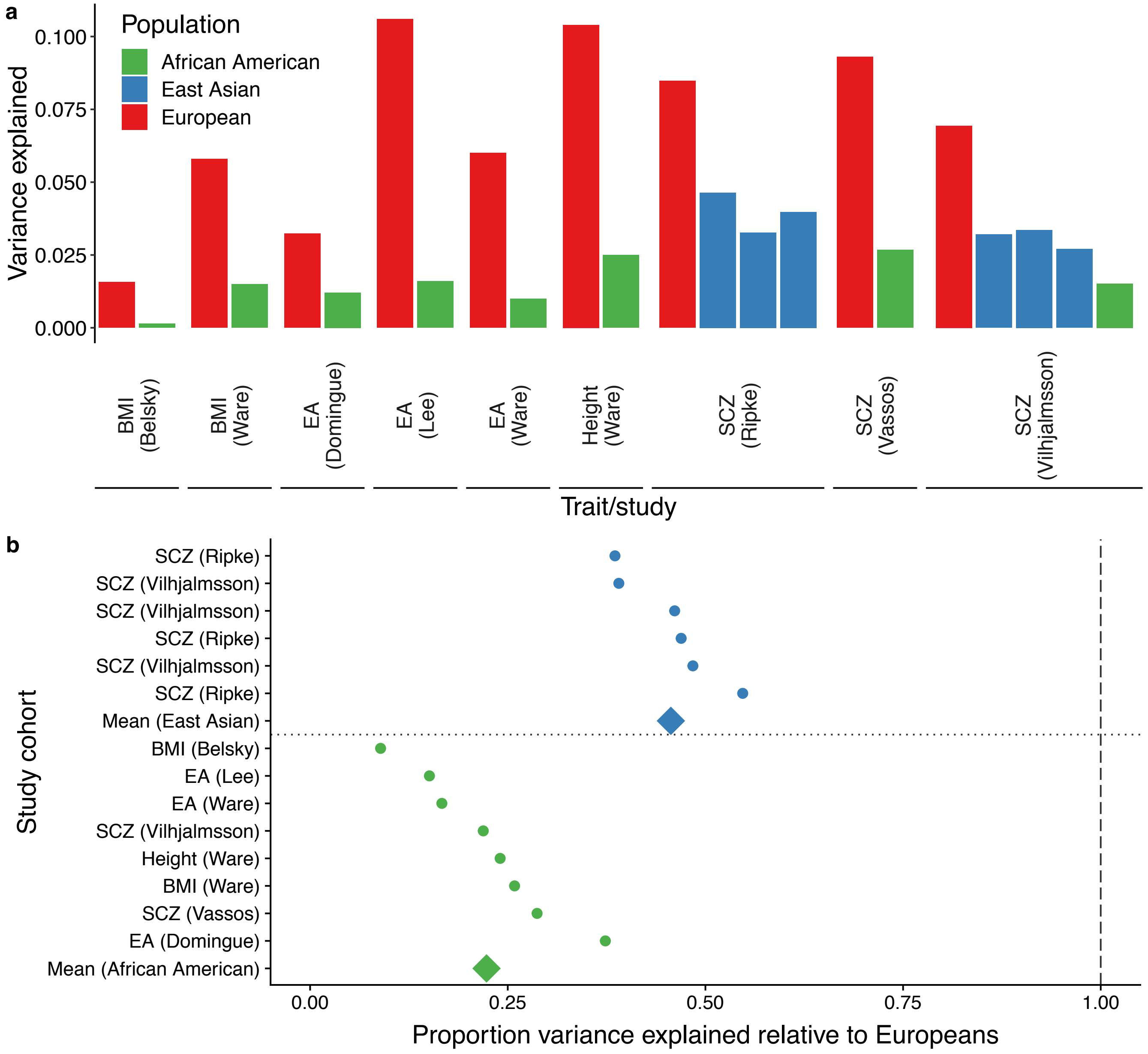


**Supplementary Figure 13. Empirical comparison of phenotypic variance explained across populations using polygenic scores computed with European GWAS**. All GWAS studies included here were conducted in European ancestry populations, with PRS calculated and evaluated in independent European, East Asian, and African American target cohorts. The European study biases result in the highest prediction accuracies in independent European cohorts, followed by declining accuracy with increased genetic divergence from Europe. **a**) Proportion of variance explained in each of the original studies. **b**) Relative proportion of variance explained in each population with respect to an independent European target population in each study. The diminished proportion of variance explained in East Asian and African American populations relative to Europeans is remarkably consistent despite differing genetic architectures, prediction methods, and accuracy metrics due to similar population histories within these cohorts. BMI = body mass index, EA = educational attainment, and SCZ = schizophrenia. Colors show the same populations in **a**) and **b**).
